# Systematic mapping of emergent transcriptional states in interacting single-cell dyads by Cell-Cell-seq

**DOI:** 10.64898/2026.02.05.704136

**Authors:** Sevana Baghdasarian, Qingyang Wang, Justin Langerman, Zhiyuan Mao, Heather Wright, Caitlin Gee, Donghui Cheng, Jami McLaughlin, John K. Lee, Xiaojing Chen, K. Christopher Garcia, Jingyi Jessica Li, Owen N. Witte, Kathrin Plath, Dino Di Carlo

## Abstract

Cell–cell interactions drive rapid and heterogeneous changes in gene expression, yet most transcriptomic methods either dissociate cells, losing pair identity and interaction timing, or infer communication indirectly from ligand–receptor co-expression. Here we present Cell-Cell-seq, a scalable workflow for profiling defined cell pairs (“dyads”) with single-cell transcriptomic resolution. Cell-Cell-seq uses cavity-containing hydrogel microparticles (Nanovials) to confine two cells, synchronize contact onset, protect fragile conjugates during handling and sorting, and interface directly with droplet-based RNA sequencing. Using antigen-matched prostate tumor cells and engineered T cells as a model system, Cell-Cell-seq captured thousands of tumor–T cell dyads and revealed broad functional and transcriptional heterogeneity across interactions. Dyads unmasked transient activation programs that were obscured in standard well-plate co-culture, consistent with asynchronous contact in bulk assays. To distinguish interaction-induced programs from the composite nature of dyad transcriptomes, we developed a pseudo-mixing framework that generates in silico pseudo-dyads to construct an empirical null distribution under “no interaction,” enabling statistically robust identification of emergent genes and partner-resolved attribution of responses. Dyad-resolved analysis further revealed coordinated cross-cell programs, including coupled chemokine expression consistent with bidirectional paracrine signaling and inverse coupling between tumor immunoregulatory programs and T cell activation. Finally, we introduce ccRepair to correct compositional dilution in mixed transcriptomes, improving interpretability while preserving genuine cross-cell coordination. Together, Cell-Cell-seq provides a generalizable platform for dissecting immune synapse biology and mapping interaction-dependent programs across heterogeneous cell populations, with applications in profiling tumor–immune communication and functionally screening immunotherapies.

## 1. Introduction

Cell–cell communication is the foundation of multicellular life. From the earliest developmental programs to the dynamics of immune defense, neural signaling, and host–microbe symbioses, cells continuously exchange information that reshapes their gene expression and behavior. These reciprocal interactions enable tissues to coordinate complex functions, while their dysregulation contributes to disorders ranging from cancer to autoimmunity and neurodegeneration. Understanding how transcriptional programs are induced by specific cellular interactions is therefore a central challenge across all of biology^1^.

Single-cell RNA sequencing (scRNA-seq) has transformed biology by providing transcriptome-wide profiles of individual cells, uncovering heterogeneity across tissues and disease states^2^. Yet, by requiring dissociation of cells into suspensions, scRNA-seq largely eliminates spatial and relational context, making it difficult to determine which cells were directly communicating^3^. Computational frameworks that infer interactions from ligand–receptor co-expression provide valuable hypotheses^4^ but remain indirect, describing potential rather than realized communication, and rarely are validated by direct analysis of interacting cell dyads. To recover this missing relational context, recent efforts have turned to spatial transcriptomics and other in situ profiling methods.

Spatial transcriptomics preserves tissue organization and provides powerful maps of gene expression across microenvironments^5^. However, most platforms still measure transcripts from regions that contain multiple cells, limiting unambiguous assignment of expression changes to a defined cell pair. Even in emerging higher-resolution approaches, transcript localization relies on computational segmentation and boundary assignment that remain imperfect, and the measured signal can reflect partial-volume contributions from neighboring cells. Moreover, adjacent cells are embedded within broader tissue microenvironments, so observed expression patterns integrate both direct contact-dependent responses and indirect influences (e.g., paracrine or longer-range signaling). As a result, spatial methods provide rich contextual snapshots of gene expression in tissues, but they are not designed to isolate the causal transcriptional programs induced specifically by a single cell–cell interaction.

To disentangle contact-dependent programs from broader microenvironmental influences, approaches are needed that capture transcriptional changes from single, physically interacting cell pairs isolated as defined units (hereafter termed dyads) and scale to hundreds to thousands of interactions to resolve response heterogeneity. Such methodologies would provide a causal framework for mapping interaction-induced programs across biological systems and offer a foundation for systematically linking defined cell-cell communication nodes to functional outcomes in health and disease.

Creating stable dyads at scale and deconvolving their mixed transcriptomes are two key challenges to study cell-cell interactions. In conventional suspension or well-based co-cultures, cells can migrate, detach, or re-partner, making it difficult to control interaction timing and to preserve fragile contacts during handling and enrichment. These limitations motivate pairing strategies that physically constrain two cells as a defined unit, reducing degrees of freedom for cell motion and enforcing sustained proximity, so that contact duration can be controlled and measured across large numbers of interactions. However, even when intact dyads are captured for sequencing, the resulting composite transcriptomes obscure which changes arise in each cell, necessitating methods that can assign interaction-induced programs to the appropriate partner cell. Guided by these constraints, we sought a pairing format that physically stabilizes dyads during handling and is directly compatible with droplet-based sequencing. We hypothesized that Nanovials, cavity-containing hydrogel microparticles, could serve as stable and protective compartments for profiling defined cell-cell interactions, while remaining compatible with droplet-based single-cell transcriptomic workflows^6^. The inner cavity confines two cells within a microscale volume, limiting lateral motion and reducing opportunities for additional cells to join the pair. The cavity surface can be functionalized to retain diverse adherent and suspension cell types in sustained proximity for controlled interaction durations^7,8^. Here, we operationally define an ‘interacting dyad’ as a two-cell pair in direct physical contact within the cavity, validated experimentally. This physical confinement also buffers fragile contacts from disruptive shear during routine handling and, when used, flow-based enrichment/sorting, preserving transient or weak interactions that would otherwise be lost. In addition, cavity-immobilized capture reagents can retain secreted proteins, enabling linkage of dyad transcriptional states to functional outputs such as cytokine secretion.

By integrating these features, we established a generalized **Cell-Cell-seq** workflow that enables systematic capture of interacting cell dyads (Figure 1). We applied this workflow to profile and investigate a previously characterized cancer–T cell interaction: a prostate cancer cell line expressing prostatic acid phosphatase (PAP) and engineered T cells with cognate T cell receptors (TCRs)^9,10^. Recognition of cancer cells by T cells is primarily initiated through TCR–pMHC binding and further facilitated by multiple surface ligand–receptor pairs as well as secreted cytokines. While patterns of T cell activation have been relatively well annotated, the heterogeneity of cancer–T cell interactions remains largely unexplored, making this model an ideal system for evaluation by Cell-Cell-seq.

**Figure 1.**
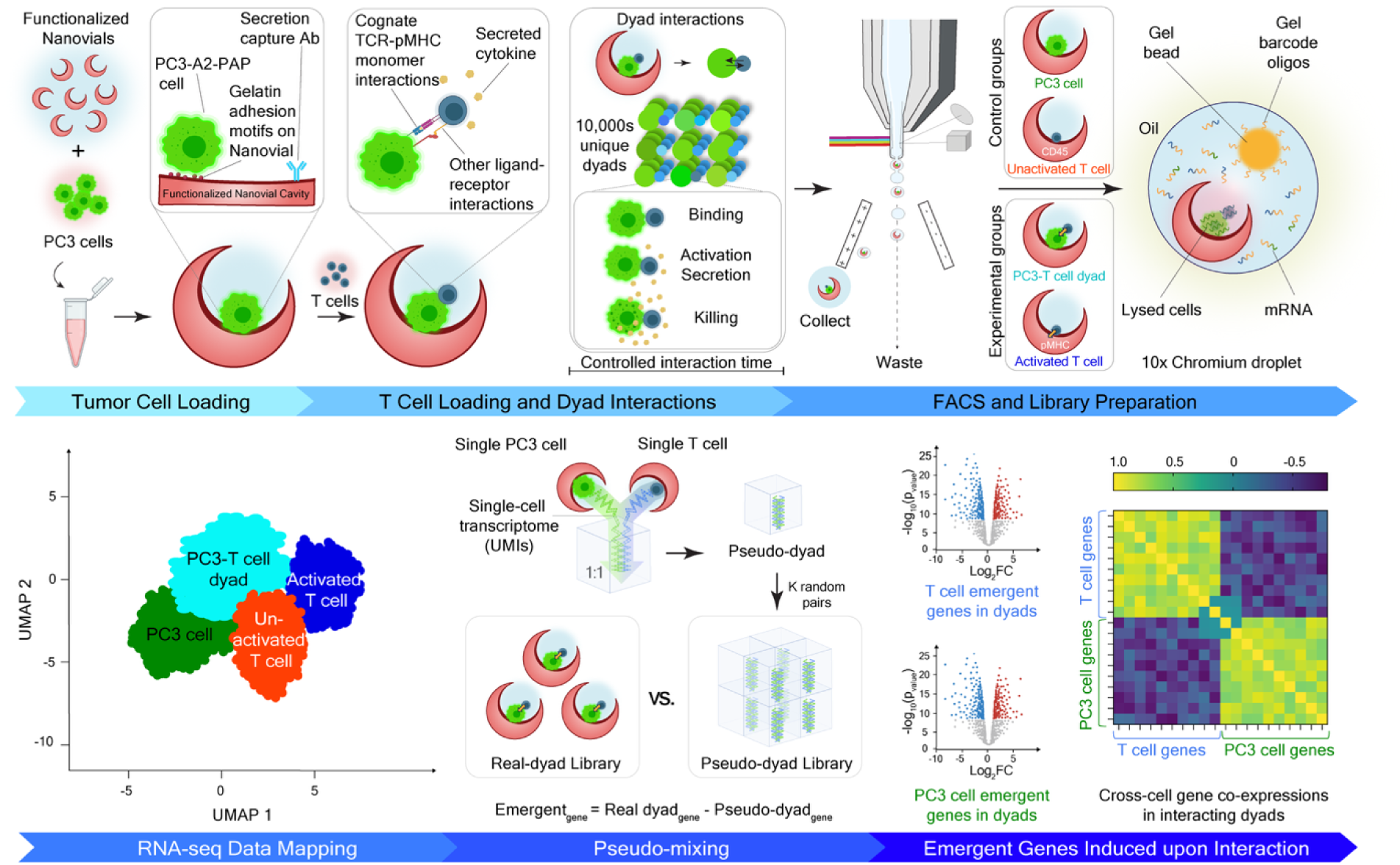
Cell-Cell-seq workflow and analysis overview. **(top row)** PC3-A2-PAP tumor cells are first loaded in Nanovials functionalized for adhesion and secretion capture, followed by addition of antigen-specific T cells to create thousands of defined tumor – T cell dyads. After a controlled interaction time, Nanovials containing interacting dyads were sorted and loaded directly into a 10x Genomics GEM-X chip to generate barcoded composite cDNA of each dyad. Control groups of cells loaded separately on Nanovials with no interaction (PC3 singlets, unactivated T cell singlets) and experimental groups (PC3-T cell dyads and pMHC-coated Nanovials with bound T cells) are sorted for enrichment and processed in parallel. **(bottom row)** Data analysis begins with UMAP visualization of major classes (PC3 singlets, unactivated T singlets, pMHC-activated T cells, and paired PC3-T cell dyads). “Pseudo-mixing” creates a library of computational pseudo-dyad controls lacking any interaction-specific gene expression by mixing gene counts from single PC3 and single T cells selected randomly. Comparing real dyads with computationally generated pseudo-dyads identifies emergent (interaction-induced) genes using differential expression testing. Following assignment to cell of origin, representative volcano plots illustrate emergent T-cell and PC3-cell genes, and a cross-cell correlation heat map highlights interaction-dependent co-expression.

To resolve the challenge of composite transcriptomes, we developed a computational framework for Cell-Cell-seq that rigorously distinguishes interaction-driven signals from background (Figure 1). First, by comparing true dyads with a library of pseudo-mixed controls generated from independently sequenced single cells, we identify genes that are uniquely perturbed upon direct interaction. Second, transcripts are assigned to each interacting cell type leveraging baseline controls, a generalizable feature for studying diverse pairings. Third, by incorporating control conditions, such as peptide major histocompatibility complex I (pMHC)-coated Nanovials that activate T cells in the absence of tumor cells, we isolate induced genes that require reciprocal communication. Together, these strategies uncover emergent gene expression networks that cannot be inferred from single-cell datasets alone. For example, Cell-Cell-seq revealed consistent negative correlations between T cell activation markers and tumor-derived immunosuppressive factors, such as TM4SF1 and DKK1, across independent experiments, highlighting the ability to identify coupled gene programs and clinically actionable axes of immune evasion. More broadly, this framework enables systematic identification of interaction-specific targets for drug discovery and provides a foundation for building predictive models of cell–cell communication in health and disease.

## 2. Results

### Nanovials enable stable co-loading of tumor and T cells

To establish a platform for studying defined cell–cell interactions, we first evaluated whether Nanovials could stably capture dyads of tumor and immune cells. PC3 prostate cancer cells engineered to express prostatic acid phosphatase (PAP) and the MHC-I allele HLA-A*02:01 (PC3-A2-PAP; dual-labeled by co-expression of GFP and RFP for fluorescence tracking) efficiently adhered to Nanovial cavities. These cells are able to present a PAP-derived peptide to antigen-specific T cells creating a stable foundation for subsequent T cell loading (Figure 2A, Supplementary Figure 1). Unless otherwise noted, we use “PC3” to refer to this engineered line (PC3– A2-P AP) throughout the manuscript.

**Figure 2.**
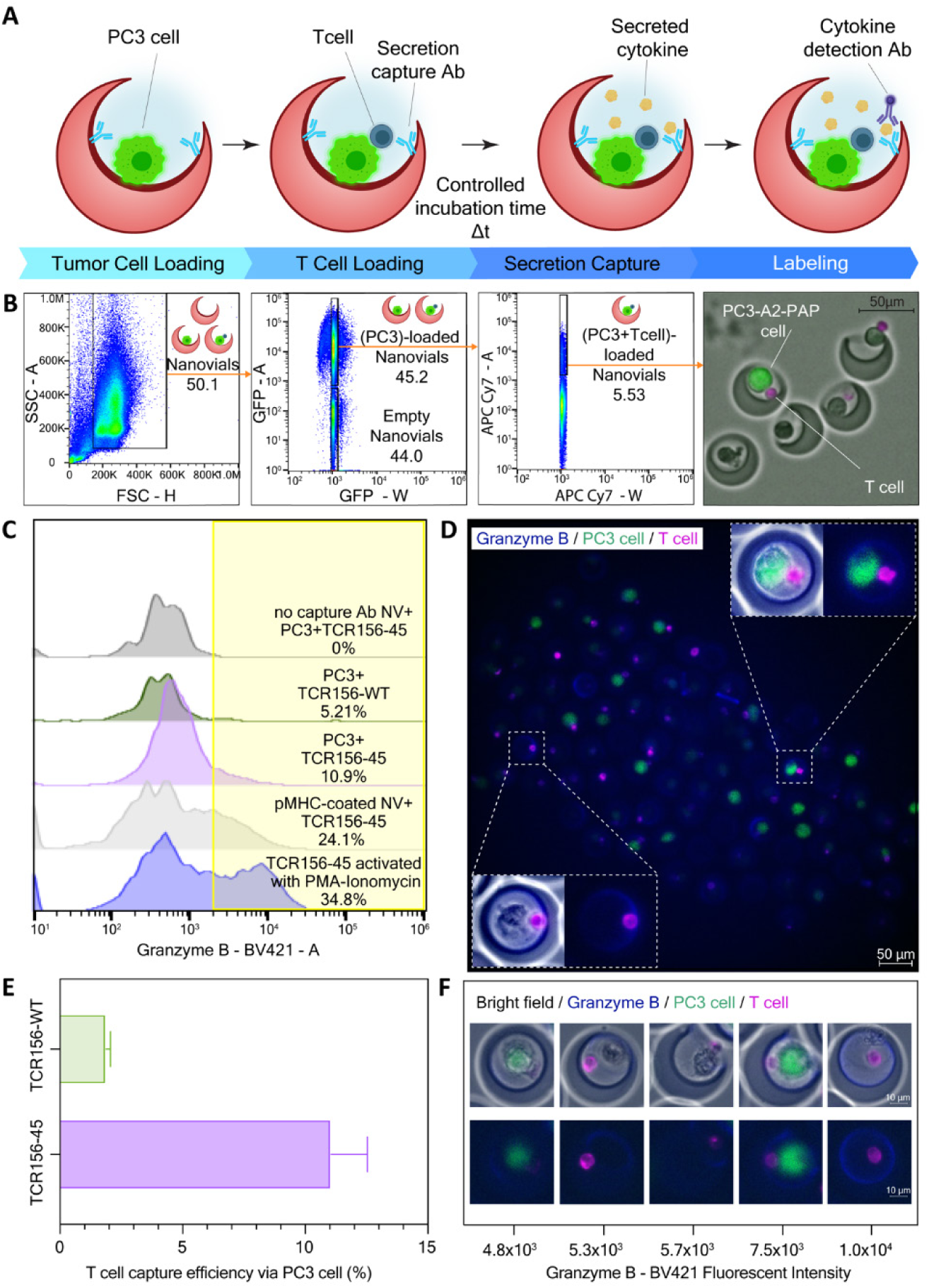
| Pairing tumor and T cells on Nanovials leads to cytotoxic responses. (**A**) Workflow schematic for the granzyme B secretion assay. PC3-A2-PAP tumor cells are first adhered inside gelatin-coated Nanovial cavities pre-functionalized with anti-granzyme B capture antibodies, T cells are added to establish defined contact for a controlled incubation interval (Δt), and secreted granzyme B bound locally is revealed with a fluorescent detection antibody. **(B)** Representative flow cytometry gates: identification of all Nanovials (FSC/SSC), enrichment of PC3-loaded Nanovials (GFP+), and detection of PC3-T cell dyads (T-cell marker, APC-Cy7). Right, bright-field/fluorescence microscopic image showing sorted Nanovials containing PC3-A2-PAP cells (green) paired with T cells (magenta). Scale bar, 50 µm. (**C**) Flow cytometry histograms of captured granzyme B signal across conditions: (i) no capture antibody (control for setting the signal threshold), (ii) PC3+TCR156-WT (weak TCR), (iii) PC3+TCR156-45 (potent TCR), (iv) pMHC-coated Nanovials+TCR156-45 (artificial antigen presentation), and (v) PMA/ionomycin activation (positive control). Percentages of events with granzyme B above the signal threshold (yellow region) are shown.

To assess whether co-capture of T cells is primarily and specifically driven by TCR–pMHC recognition, we used two previously developed TCRs^10^, TCR156-WT and TCR156-45, both targeting the same PAP epitope (TLMSAMTNL) presented by HLA-A*02:01 but differing in their dwell time on pMHCs upon binding. These TCRs are structurally highly similar, differing by only two amino acids. In TCR156-45, selected mutations were created to introduce “catch bond” interactions that prolong dissociation, thereby stabilizing cancer–T cell interactions. Prior studies showed that T cells expressing TCR156-45 exhibit strong cytotoxicity against PC3 cells, whereas those expressing TCR156-WT do not.

Brightfield and fluorescence microscopy confirmed stable confinement of a PC3 cell together with a T cell within individual Nanovial cavities (Figure 2B, D), with both cell types fitting within the cavity while maintaining direct cell–cell contact. We next asked whether co-capture reflected the biophysical strength of the cognate pMHC–TCR interaction. T cells engineered with TCR156-45 bound to PC3-loaded Nanovials with higher efficiency than T cells expressing the wild-type receptor (Figure 2E), consistent with preferential stabilization of antigen-specific contacts. Finally, flow cytometry enabled robust enrichment of double-positive Nanovials containing both PC3 and T cells (Figure 2B), supporting scalable recovery of defined dyads for downstream handling and analysis.

### Functional tumor–T cell interactions captured within Nanovials

We next asked whether co-loading preserved functional responses associated with tumor cell recognition by T cells. To quantify productive cytotoxic activation, we functionalized Nanovials with granzyme B capture antibodies (Figure 2A) and measured accumulated granzyme B secretion from individual tumor–T cell dyads. Fluorescence microscopy confirmed granzyme B capture on a subset of TCR156-45 dyads with PC3 cells (Figure 2C,D), and across hundreds of dyads granzyme B signal spanned more than an order of magnitude, revealing marked functional heterogeneity (Figure 2F). Approximately 11% of tumor–T cell dyads containing TCR156-45 T cells exceeded the granzyme B secretion threshold, compared with ∼5% for TCR156-WT dyads and <1% in no-capture-antibody controls (Figure 2C). As positive benchmarks, pMHC-coated Nanovials and PMA/ionomycin stimulation yielded 24% and 39% of dyads above threshold, respectively (Figure 2C), supporting that granzyme B capture reports productive T cell activation rather than nonspecific retention. Granzyme B accumulation increased with interaction time (Supplementary Figure 2): in antigen-matched TCR156-45 dyads, the fraction above threshold rose from ∼0.7% at 3 h to ∼15% at 7 h and ∼21% at 11 h, whereas TCR156-WT dyads remained near baseline (∼1–2%) over the same interval. Together, these results show that Nanovials preserve tumor–T cell dyads and enable functional readouts of activation-dependent effector responses.

### Integration with single-cell transcriptomics

To link defined cell-cell interactions with transcriptional state, we introduced sorted Nanovials containing tumor–T cell dyads into a droplet microfluidics scRNA-seq workflow (10x Genomics Chromium GEM-X). Because epithelial tumor cells are substantially larger than T cells and typically contain more total RNA, we empirically optimized Nanovial dimensions to (i) reliably accommodate a tumor–T cell dyad within the cavity and (ii) maintain robust emulsion generation without clogging, while also understanding the influence of the more RNA-rich partner on dyad libraries (Supplementary Figure 3). In parallel, we profiled matched PC3-only and T cell-only singlets (including activated T cell controls) to provide reference states and to build an additive ‘pseudo-dyad’ null model used later to identify interaction-induced programs. We successfully obtained transcriptomes from dyads, as well as from PC3 cells alone, T cells bound via anti-CD45 without tumor exposure, and T cells activated on pMHC-coated Nanovials (Supplementary Figure 4). To focus on early transcriptional responses while maintaining tumor cell viability and intact transcriptomes, dyad interactions were limited to 3 hours prior to enrichment and library preparation.

### Interaction-specific transcriptional states and heterogeneity

Across all Nanovial events, a UMAP embedding separated PC3 singlets, unactivated T cell singlets, pMHC-activated T cell controls, and PC3–T cell dyads into distinct clusters (Figure 3A). Consistent with this interpretation, dyads showed intermediate expression of canonical lineage markers (T cell: *PTPRC*, *TRAC*; PC3: *EPCAM*, *HOXB13*), while a subset of genes was selectively elevated in dyads relative to either parent population (e.g., *IRF1*, *TNF*) (Figure 3B).

**Figure 3.**
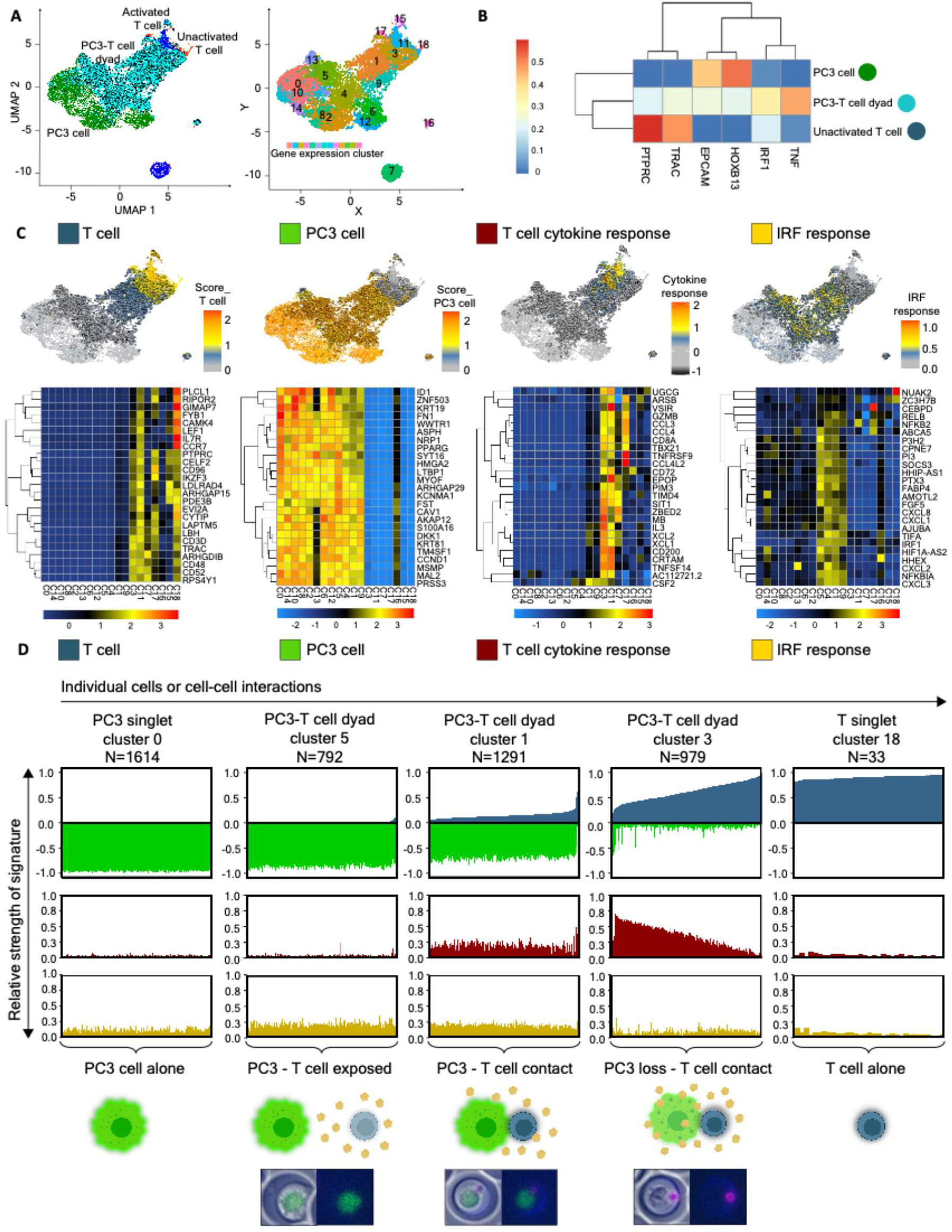
Interaction-dependent gene programs revealed in tumor-T cell dyads. (**A**) UMAP embedding of all Nanovial events (left) and corresponding cluster labels (right). Points correspond to transcriptomes recovered from Nanovials, including singlets and dyads. **(B)** Heat map of mean gene expression for representative T cell genes (*PTRC*, *TRAC*), PC3 cell genes (*EPCAM*, *HOXB13*), and induced genes in interacting cells (*IRF1*, *TNF*) across PC3 cell, unactivated T cell, and dyad (composite) populations. **(C)** Gene module-score overlays and heat maps for canonical T cell, PC3 cell, T cell cytokine response, and interferon response (IRF) signatures showing induced responses in activated T cells and dyads. Top genes representing each signature are shown in heat maps (mean expression by cluster/condition, as indicated). **(D)** Ordering cells by relative gene signature strength across clusters (0, 5, 1, 3, 18) shows graded responses across interaction states. Within each column, the top track shows relative PC3 (green) versus T cell (blue) signature strength, with cytokine-response (red) and IRF-response (yellow) scores shown below; events are ordered left-to-right by relative signature balance to illustrate graded interaction states. Clusters 5, 1, and 3 for example represent distinct interaction states reflecting cell-cell interaction heterogeneity. Representative images beneath each column illustrate the corresponding contexts consistent with the gene expression data.

To interpret these dyad-associated responses, we overlaid gene-module scores corresponding to canonical T cell and PC3 signatures, as well as an interferon response (IRF) program and a T cell cytokine/effector program (Figure 3C). IRF and T cell cytokine/effector programs reflected a subset of gene modules most strongly associated with dyad responses (Supplementary Figure 5). Module-score overlays revealed substantial dyad-to-dyad heterogeneity: some dyads strongly induced both IRF and cytokine/effector programs, others preferentially induced one program, and a minority showed minimal induction (Figure 3C). The cytokine/effector program observed in dyads partially overlapped with the transcriptional program in pMHC-activated T cell controls (Figure 3C), and within dyads, memory-associated genes (e.g., *IL7R*, *CCR7*) were reduced while activation/effector genes (e.g., *GZMB*, *TNFRSF9*/CD137) were increased, consistent with emergence of an activated T cell phenotype (Figure 3C).

Notably, interaction timing was held constant (1.5 h loading and 3 h co-culture), yet dyads occupied a continuum of composite states. Figure 3D summarizes dyad-associated clusters by ordering events along a continuum of relative cell-type signature strength and response programs. In the leftmost state (PC-3 singlets; cluster 0), transcriptomes are dominated by the PC3 signature (green) with minimal T cell contribution (blue). In dyad clusters, events span graded composite states: cluster 5 (“PC3–T cell exposed”) shows modest T cell signal with limited cytokine induction; cluster 1 (“PC3–T cell contact”) exhibits stronger T cell contribution accompanied by increased cytokine-response scoring (red); and cluster 3 (“PC3 loss–T cell contact”) is characterized by high cytokine-response scores alongside markedly reduced PC3 signature. The rightmost state (T cell singlets; cluster 18) is dominated by the T cell signature with minimal PC3 contribution. Across these states, IRF-response scoring (yellow) is elevated in dyad states and varies independently of the cytokine program; representative images aligned to each column illustrate these corresponding Nanovial contexts (Figure 3D). To test robustness, we performed an independent Cell-Cell-seq run and observed similar dyad-associated subpopulations (Supplementary Figure 6). In parallel, bulk co-culture captured a broad interferon response in PC3 cells but did not resolve the coordinated high-IRF/high-cytokine dyad states or the tumor-low/T cell–dominant endpoint states, consistent with averaging and loss of pair identity in standard workflows (Supplementary Figure 6). Because each dyad transcriptome is a composite mixture of tumor and T cell RNA, identifying interaction-induced changes requires a statistically grounded baseline for expression under ‘no interaction’ that reflects realistic variability in both partners. We therefore developed a computational pseudo-mixing framework that constructs an empirical null distribution from a large library of in silico pseudo-dyads, enabling quantitative testing of interaction-dependent gene programs.

### Pseudo-mixing identifies emergent genes induced upon interaction

A central challenge in analyzing transcriptomes of cell dyads is distinguishing true interaction-induced programs from the background signal that arises simply from combining two independent cell transcriptomes. A naïve approach, subtracting an “average” PC3 profile plus an “average” T cell profile, fails because both cell types exhibit substantial cell-to-cell variability (and, for tumor cells in particular, broad transcriptional dispersion), such that many genes can appear “changed” simply due to which two cells happened to be paired.

To address this, we developed a computational pseudo-mixing framework (Supplementary Figure 7) that constructs an empirical null distribution for mixed dyad transcriptomes by generating a large library of pseudo-dyads. Specifically, we randomly sampled transcriptomes from non-interacting PC3 cells and non-interacting T cells and summed them to form many *in silico* dyads spanning the full range of plausible PC3×T transcriptional combinations (Figure 4A, Supplementary Figure 7). This strategy is conceptually related to approaches used to model technical artifacts such as scRNA-seq doublets, where synthetic mixtures are generated to represent expected mixed profiles^11^. However, rather than using pseudo-mixtures solely to flag artifacts, we purposefully use them to define the expected distribution of expression for each gene under “no interaction”, conditioned on realistic variability in both partners.

**Figure 4.**
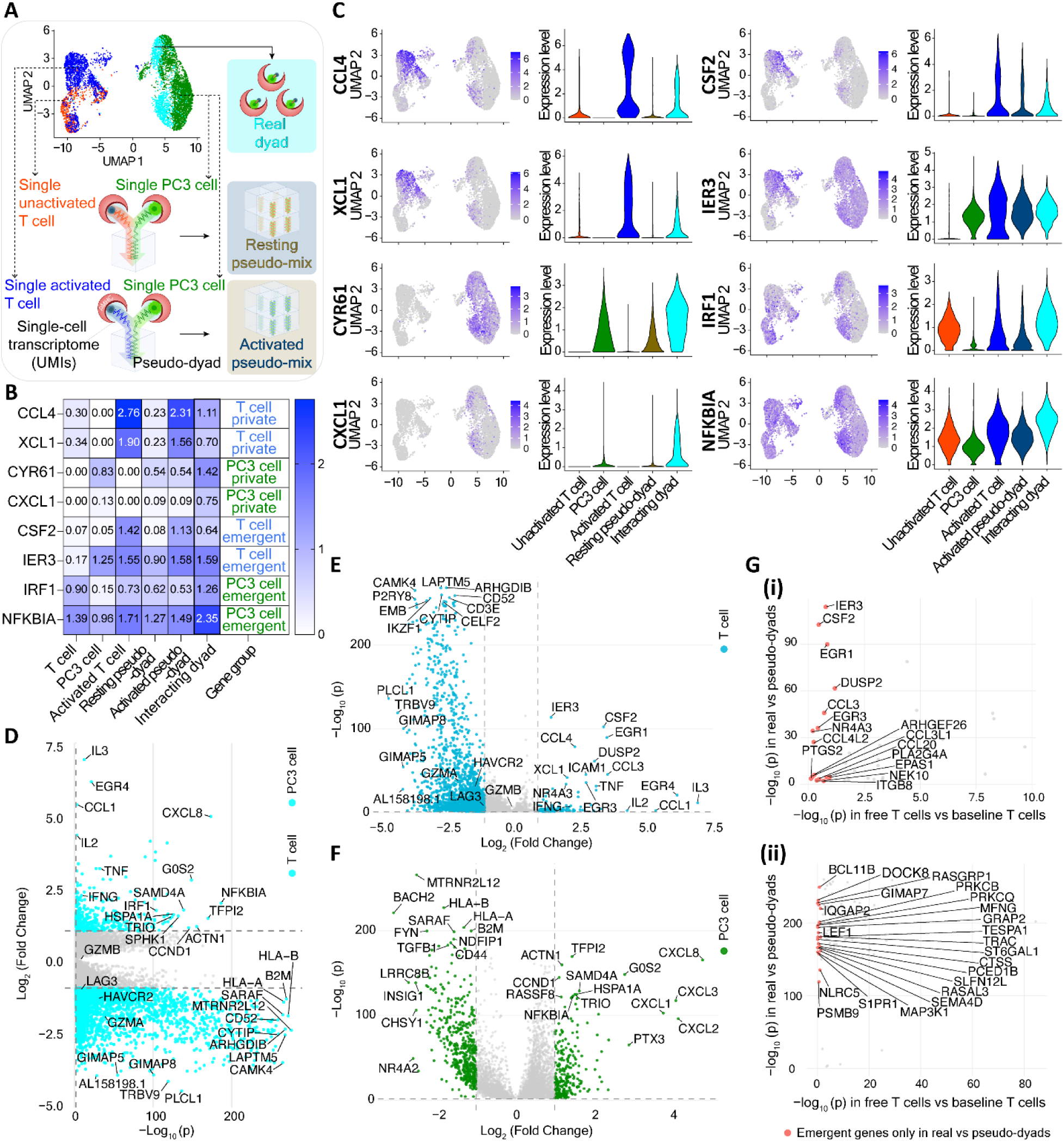
Interaction-induced emergent gene expression and cell assignment. (**A**) Pipeline design for comparing real dyads to pseudo-dyads. UMIs from randomly selected single PC3 and single T cells are mixed to build baseline pseudo-dyad gene expression libraries that lack any interaction-induced gene expression. This is repeated separately for resting T cells to create resting pseudo-dyad libraries and activated T cells to build activated pseudo-dyad libraries. (**B**) Table illustrating the assignment process for cell-specific gene expression. Private genes for each cell type are based on no baseline expression in the other cell type. Emergent genes are assigned based on a statistical test against the activated pseudo-dyad library. (**C**) UMAP feature plots and violin plots for representative genes from each attribution category: private T cell genes (e.g., *CCL4*, *XCL1*), private PC3 genes (e.g., *CXCL1*, *CYR61*), and shared/interaction-induced genes (e.g., *CSF2*, *IER3*, *IRF1*, *NFKBIA*). Violin plots show per-cell expression distributions stratified by condition. Pseudo-dyads are not included in the UMAP embedding. (**D**) Volcano plots for all differentially expressed genes when comparing between real dyads and resting pseudo-dyad libraries. (**E**) T cell assigned genes. (**F**) PC3 cell assigned genes. **(G)** p-value rankings for **(i)** upregulated T cell assigned genes and **(ii)** downregulated T cell assigned genes, comparing T cells co-cultured with PC3 cells in well plates (x-axis) vs. controlled pairing in Nanovials (y-axis). Immediate early genes are labeled and appear prominently along the y-axis, reflecting the synchronized dyad co-culture in Nanovials.

These pseudo-dyads therefore provide a rigorous baseline for what a mixed transcriptome should look like in the absence of communication. By comparing each observed dyad to this library (rather than to a single composite expectation), we identify emergent gene expression as genes whose expression in true dyads deviates significantly from the pseudo-dyad distribution—i.e., changes that cannot be explained by random pairing of independent cells and instead reflect transcriptional programs induced by direct interaction.

To attribute emergent genes to the cell-of-origin within the dyad (tumor versus T cell), we implemented two complementary strategies. First, **private gene mapping** leveraged transcripts detectably expressed in only one parent population under non-interacting reference conditions. For example, *CCL4* and *XCL1* were observed in T cells but not PC3 cells, while *CXCL1* and *CYR61* were restricted to PC3 cells (Figure 4B). Emergent expression of these genes in dyads was therefore assigned to the most likely originating partner. Because interaction could, in principle, induce low-level expression of some transcripts in the opposite cell type, this ‘private gene’ criterion is intentionally conservative and primarily supports assignment for strongly lineage-biased genes.

We next expanded gene assignment using **activated-state pseudo-mixing** (Supplementary Figure 7), which distinguishes dyad-specific programs from transcriptional changes expected from T cell activation alone. Specifically, we compared emergent genes in true dyads to pseudo-dyads generated from non-interacting PC3 cells paired *in silico* with T cells activated on cognate PAP pMHC-coated Nanovials (Figure 4B-C). Genes induced in true dyads but absent from these activated-state pseudo-dyads were attributed to PC3-specific interaction responses (e.g., *IRF1*, *NFKBIA*), whereas genes amplified in both were attributed to the T cell activation program (e.g. *CSF2* and *IER3*). Together these approaches expanded the assignable gene set to include context-dependent responses such as *CSF2*, *IER3*, and *NR4A3* in T cells, and stress-associated genes in PC3 cells (Figure 4B-C).

Having established these gene-attribution categories (private T cell, private PC3, and shared/interaction-induced), we next visualized how each class localizes across the joint embedding of singlets, activated controls, and dyads. UMAP feature plots and associated violin plots (Figure 4C) revealed distinct enrichment patterns for genes in each attribution category. UMAP overlays illustrate expression across experimentally measured singlet and dyad transcriptomes, whereas pseudo-dyad states are included in the violin plots only; violins quantify expression across unactivated T cell, activated T cell, PC3 cell, resting pseudo-dyad, activated pseudo-dyad, and interacting dyad conditions. Private T cell–biased genes such as *CCL4* and *XCL1* were highly expressed in subpopulations of activated T cells and interacting dyads but were not detected in PC3 singlets. Conversely, *CXCL1* and *CYR61* showed low basal expression in PC3 cells that was strongly increased in a subset of dyads. In contrast, shared or interaction-induced genes such as *IER3*, *IRF1*, and *NFKBIA* were low in resting pseudo-dyads but became selectively concentrated within the dyad-associated cluster, consistent with programs induced by cell–cell engagement. Comparing activated pseudo-dyads to true interacting dyads further distinguished tumor-sided interaction responses from activation-linked genes: *IRF1* remained substantially lower in activated pseudo-dyads than in interacting dyads, whereas *IER3* exhibited similar distributions in both conditions, suggesting a predominantly T cell–sided response rather than PC3 induction.

We next performed genome-wide differential expression comparing measured dyads to computational pseudo-dyads to define an empirical-null-based set of interaction-associated (‘emergent’) genes (Figure 4D). Whereas Figure 4C highlights representative examples, this analysis systematically identifies transcripts whose expression deviates from the distribution expected under additive mixing of independently profiled PC3 and T cell transcriptomes. Across two independent Cell-Cell-seq experiments (runs 1 and 2), we observed a reproducible emergent gene set, with a larger fraction of significant genes downregulated in dyads relative to pseudo-dyads than upregulated (Figure 4D). Concordance across runs was supported by overlap among the most significant emergent genes and by correlation of per-gene effect sizes (Pearson r=0.63; Figure 4D). Importantly, the number and identity of emergent genes were robust to reasonable changes in pseudo-dyad mixing weights (e.g., equal versus PC3-weighted mixtures; Supplementary Figure 8). Together, these results show that pseudo-mixing provides a statistically grounded baseline for isolating interaction-associated transcriptional signals, which we next assign to the tumor or T cell compartment.

To further resolve the cellular origin of emergent transcriptional programs, we applied our deconvolution framework to assign the larger set of emergent dyad genes to either the T cell or PC3 compartment, yielding compartment-specific volcano plots (Figure 4E,F). Representative downregulated transcripts included *BCL11B*, *GIMAP5-8*^12^, and *CXCR4*^13^, while upregulated transcripts included immediate-early and activation-associated genes such as *IER3*^14^, *CCL4*^15^, and *EGR1*^16,17^ (Figure 4E). Notably, *BCL11B* has been reported to decrease upon engineered T cell activation and its perturbation has been linked to enhanced anti-tumor activity in adoptive T cell contexts^18,19^.

For PC3-assigned genes, upregulated transcripts were enriched for inflammatory/stress-adaptive programs and included chemokines implicated in tumor–immune crosstalk (e.g., *CXCL1/2/3/8*), whereas downregulated genes were enriched for cytokine-related signaling pathways (Figure 4F). Together, these compartment-resolved programs align with known biology: T cell activation and effector signaling alongside tumor inflammatory and stress responses, supporting the robustness of deconvolution-based gene attribution. Because Cell-Cell-seq profiles defined dyads at early time points, these signatures emphasize immediate-early and contact-proximal responses that may be obscured or remodeled in longer-term co-culture or in vivo settings.

### Early and transient interaction programs are enriched in defined dyads

To test whether defined, time-synchronized dyads reveal early and transient transcriptional responses that are diluted in conventional assays, we compared emergent gene signatures from Nanovial-confined interactions to those measured in standard well-plate bulk co-culture (effector to target ratio, E:T of 1:1). While some overlap was observed, immediate-early response genes (e.g., *IER3*, *EGR1*, *EGR3*, *NR4A3*) were among the most strongly enriched in the Cell-Cell-seq condition (Figure 4G(i)). These transcripts are rapidly and transiently induced upon stimulation, consistent with Cell-Cell-seq preserving defined pairing and synchronized interaction timing. In contrast, these signals were reduced or absent in well-plate population-averaged co-culture, where interaction onset is asynchronous and pairwise resolution is lost. Additional transcripts were downregulated upon interaction and were weakly reflected in bulk co-culture (Figure 4G(ii)), suggesting further interaction-associated programs that are obscured by population averaging. Together, these analyses show that Cell-Cell-seq enriches early, contact-proximal transcriptional programs by maintaining defined dyads under controlled timing, while the pseudo-mixing framework provides a statistically grounded baseline to identify emergent genes, assign them to the appropriate cellular source, and compare interaction programs across experimental contexts.

### Reciprocal correlations reveal coordinated cell-cell interaction programs

Before examining gene–gene relationships, we asked whether tumor–T cell dyads exhibit coordinated responses at the level of transcriptional programs. We scored each dyad for the same gene modules defined in Figure 3 (T cell signature, PC3 signature, T cell activation/cytokine response and IRF response) and compared module activities across thousands of dyads (Figure 5A–C). At this program level, the strongest positive relationship was observed between the T cell cytokine-response module and the IRF-response module (Pearson r = 0.36; Figure 5D), consistent with coordinated immune activation and tumor-associated interferon signaling in a subset of dyads. In contrast, cell-type–intrinsic modules showed weak negative correlations (e.g., T cell vs PC3 signature; r = −0.11), consistent with compositional tradeoffs in composite dyad transcriptomes. Guided by these program-level relationships, we next quantified pairwise gene–gene correlations across dyads after assigning transcript origin to the T cell or PC3 compartment using private-gene mapping (Figure 5E). This analysis revealed coordinated intercellular gene programs that were not apparent in pseudo-mixed controls (Supplementary Figure 9). Among the most prominent features was a coordinated chemokine axis, marked by positive correlations between T cell–derived chemokines (e.g., *CCL3/4*, *XCL1/2*) and tumor-derived chemokines (e.g., *CXCL1/3/8*), consistent with synchronized paracrine signaling across the immune synapse. Similar intercellular correlation structure was observed across independent runs. In contrast, a distinct block of negatively correlated gene pairs emerged, including a robust inverse relationship between the tumor-associated gene *TM4SF1* and multiple T cell activation genes (*CD3D*, *CD3E*, *CD3G*, *CD2*, *CYBA*, *CD53*)^20–23^, a signature that was absent in pseudo-dyads (Supplementary Figure 9). To further validate this relationship, we stratified dyads by *CD3E* expression and examined *TM4SF1* levels. Violin plots confirmed that dyads with high *CD3E* expression were associated with significantly lower *TM4SF1* expression in the tumor compartment (Figure 5F), consistent with stronger T cell activation programs. This coupling was not observed in pseudo-mixed controls (Figure 5G), indicating that it cannot be explained by additive mixing alone. Notably, these dyads were not confined to a single UMAP cluster (Figure 5H), suggesting that intercellular communication states span a continuum that is not fully captured by discrete clustering.

**Figure 5.**
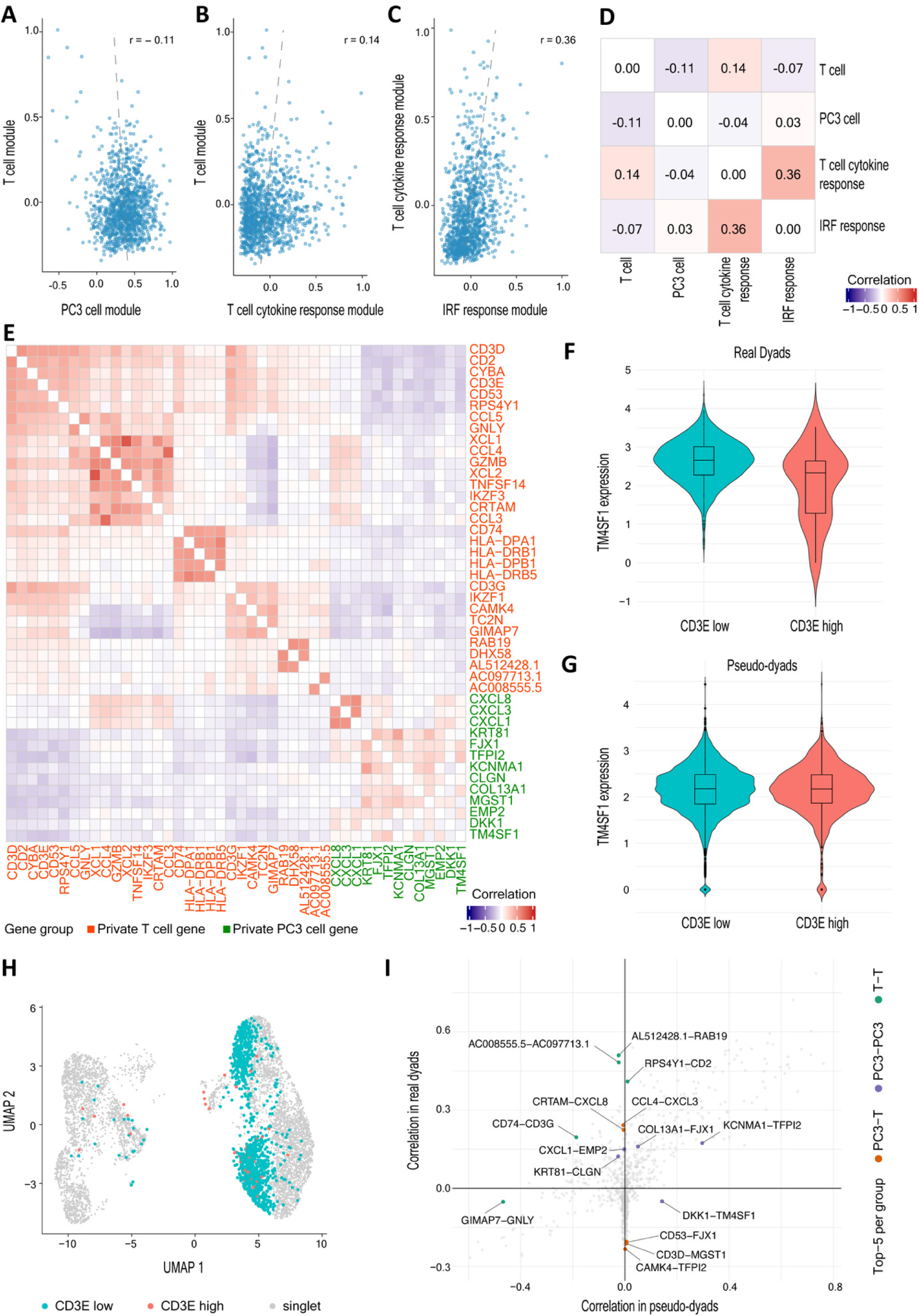
Identifying gene expression programs across interacting dyads. (**A–C**) Scatter plots of module scores for individual dyads show a weak anti-correlation between tumor and T-cell identity modules (A, Pearson’s correlation, r=-0.11), a modest positive relationship within T-cell identity and activation modules (B, Pearson’s correlation, r=0.14), and a stronger coupling between T-cell cytokine and interferon-response modules (C, Pearson’s correlation, r=0.36). (**D**) Correlation matrix of module scores across dyads (Pearson r) summarizes these trends. **(E)** Gene–gene correlation heat map for dyads is shown for genes with the largest magnitudes of Pearson’s correlation coefficient (r). Private genes assigned to T cell private (red) and PC3 cell private (green) are labeled; blocks of positive within-cell correlations and cross-cell anti-correlations are observed. (**F**) Violin plot for the gene expression distribution for an example of negative correlation (*TM4SF1* vs. *CD3E*). The *TM4SF1* distribution is shown for dyads with high *CD3E* (top 5%) and bottom 95%. The p-value when comparing the means between these distributions. (**G**) Violin plot for the gene expression distribution for the same genes in the pseudo-dyad library and corresponding p-value. (**H**) UMAP of interacting dyads with *CD3E*-high (top 5%) and *CD3E*-low (bottom 95%) groups overlaid, illustrating where these dyads reside in transcriptional space. (**I**) Gene x gene correlations plotted for real interacting vs. pseudo-dyads. The top five correlations per group comprising genes both assigned to T cells, both assigned to PC3 cells, or across T cells and PC3 cells are highlighted.

To globally assess which gene–gene correlations are specifically associated with cell–cell interaction, we compared pairwise correlation coefficients measured in real dyads (y-axis) to those in pseudo-dyads (x-axis) (Figure 5I). This analysis stratifies gene pairs into three regimes: (i) interaction-associated correlations that are absent or weak in pseudo-dyads but strong in real dyads (e.g., *CCL4*–*CXCL3*, *XCL1*–*CXCL8*), which lie high along the y-axis; (ii) interaction-associated negative correlations that are strengthened in real dyads (e.g., *CD3E*–*TM4SF1*) which fall low on the y-axis; and (iii) correlations that are similar in real and pseudo conditions, clustering near the y = x diagonal. Together, this landscape highlights coordinated intercellular gene programs that emerge during defined tumor–T cell engagement and are not explained by additive mixing alone.

### Correcting gene expression bias from mixed transcriptomes

A key challenge in analyzing dyad RNA-seq is compositional dilution: the measured transcriptome is a mixture of two cells, and when one partner contributes more RNA (or is more transcriptionally active), it can mask changes in the other. In our data, this manifests as a negative correlation between overall T cell and PC3 signature strength across dyads (Figure 5A), indicating that variation in partner RNA contribution can compress cell-type-specific signals and complicate interpretation of interaction-induced programs. Without correction, dyad-to-dyad differences can reflect how much RNA each partner contributed rather than how each partner responded to interaction, obscuring true interaction-dependent programs. Notably, the pseudo-mixing framework used to define emergent genes compares true dyads to composition-matched pseudo-dyads and is therefore largely robust to this effect; ccRepair is introduced here to improve partner-resolved interpretation and state separation across dyads.

To mitigate the compositional dilution effect, we developed ccRepair, a normalization strategy that restores each partner’s gene expression scale toward singlet-derived baselines. ccRepair (i) identifies stable “anchor” genes for T cells and PC3 from non-interacting singlets, (ii) uses these anchors to estimate the expected contribution of each cell type in an unmixed state, and (iii) rescales cell-type compartment genes within each dyad to match these expectations (Figure 6A). Conceptually, ccRepair does not “create” new expression; it corrects for unequal mixing so that biological differences among dyads are not dominated by fluctuating RNA contribution.

**Figure 6.**
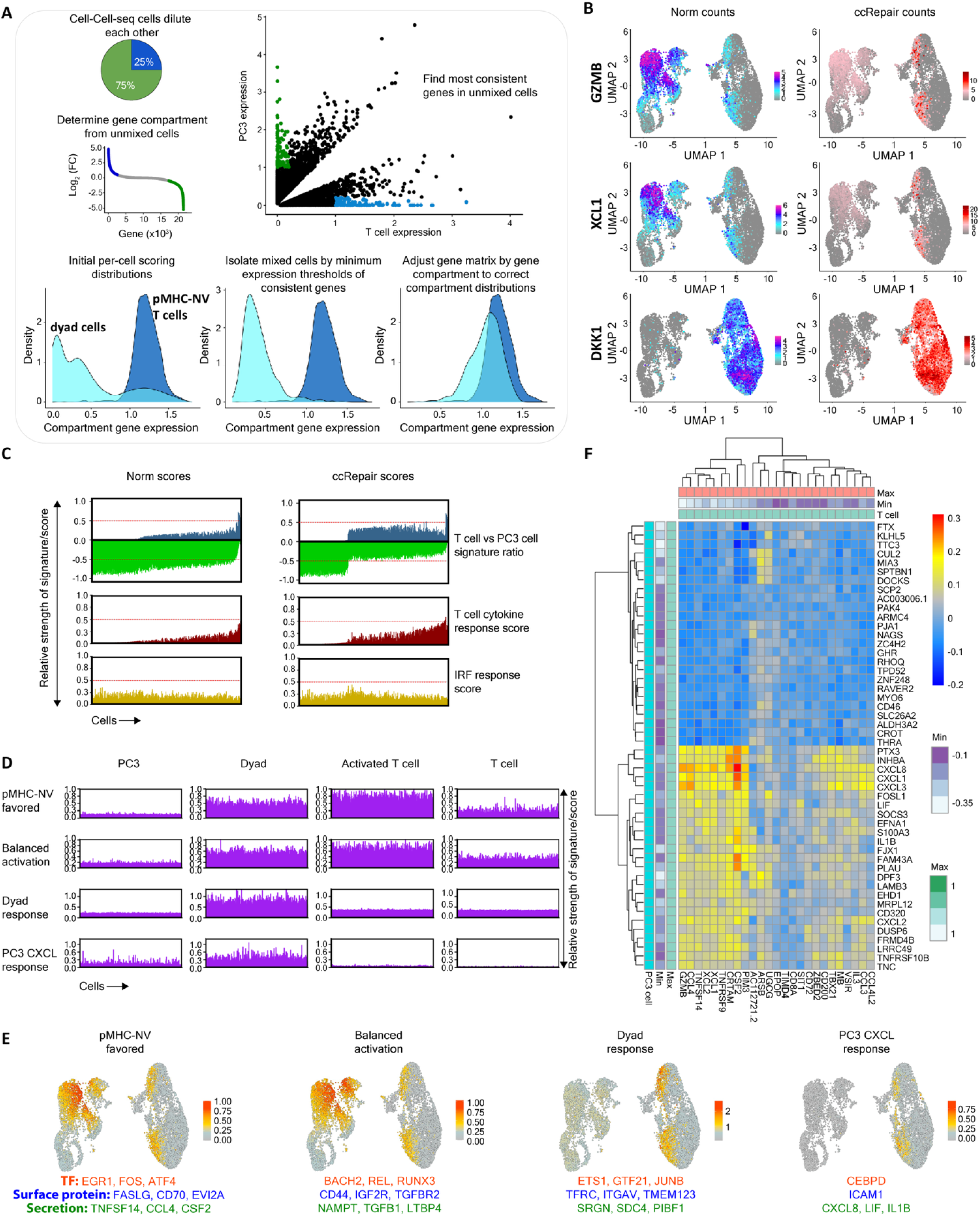
ccRepair corrects mixed-transcript dilution revealing interaction programs. (**A**) Concept: expression in dyads is a mixture of two cell types; ccRepair identifies highly consistent origin-specific genes from unmixed cells and rescales dyad counts by a cell-specific consistency ratio. **(B)** UMAPs of representative genes (*GZMB*, *XCL1*, *DKK1*) before and after ccRepair. After correction, signal from over-represented PC3 cells in dyads diminishes while it increases for the under-represented T cells in dyads. (**C**) Signature scores before/after correction show restoration of dynamic range for T cell, PC3, T cell cytokine response, and IRF response signatures in dyads. **(D)** Rank-ordered signature scores after ccRepair for four interaction modules: pMHC-NV-favored, balanced activation, dyad response, and PC3-CXCL response displayed across PC3 singlets, real dyads, activated T cell singlets, and resting T cell singlets. **(E)** UMAP overlays of the same four signature scores. For each panel, representative top gene drivers are listed beneath the map (top: transcription factors; middle: cell surface proteins; bottom: secreted mediators). (**F**) ccRepair-corrected correlation heat map of T cell cytokine response genes (x-axis) and the high and low correlated PC3-assigned genes (y-axis), and clustered by similarity.

This correction yields practical improvements. First, cell-type markers and functional genes that were attenuated by dilution become sharply resolved, including T cell effector genes (e.g., *GZMB*, *XCL1*) and tumor immunoregulatory genes (e.g., *DKK1*), enabling clearer attribution of programs to the appropriate partner (Figure 6B–C). Second, ccRepair enables cleaner program-level comparisons across conditions: using corrected signatures, we distinguish programs favored by pMHC-only activation from programs preferentially enriched in real dyads, including a PC3-specific dyad response containing adhesion/synapse-associated genes (e.g., *ICAM1*) that is not recapitulated by antigen stimulation alone (Figure 6D–E, Supplementary Figure 10).

Importantly, ccRepair preserves genuine cross-cell coordination while reducing technical distortion. After correction, coordinated multicellular modules, such as the chemokine bloom signature linking PC3 chemokines (*CXCL1/3/8*) with T cell inflammatory/effector genes (*XCL1/2, CSF2*), remain evident and become more robustly detectable (Figure 6F). Together, ccRepair improves interpretability of dyad transcriptomes by separating biology (interaction state) from mixing (RNA contribution), thereby exposing reciprocal, partner-dependent programs that would otherwise be diluted or obscured.

## 3. Discussion

Understanding how individual cells communicate in multicellular environments remains a central challenge across the life sciences from immunology and oncology to developmental biology. Here, we introduce Cell-Cell-seq, a scalable platform that physically encapsulates and transcriptionally profiles defined interacting cell dyads in Nanovial hydrogel cavities. By preserving dyad identity, controlling the timing and duration of contact, and enabling downstream multiomic readouts, Cell-Cell-seq overcomes key limitations of both dissociative single-cell methods and conventional co-culture systems. This platform provides a unique lens into the transcriptional and functional consequences of direct cell–cell interaction, capturing emergent, contact-dependent programs with single-dyad resolution.

Unlike ligand–receptor inference frameworks (e.g., CellPhoneDB^24^ and CellChat^4^), which infer putative communication by pairing the expression of ligands in one cell population with cognate receptors in another, Cell-Cell-seq directly measures the transcriptional consequences of realized cell–cell interactions and links these states to functional outcomes. Ligand–receptor methods are well-suited for constructing tissue-scale interaction networks and prioritizing candidate signaling axes, but because they operate on expression snapshots cannot on their own establish whether two cells were interacting, over what time period, or whether ligand–receptor co-expression produced a downstream response in either partner. Cell-Cell-seq is conceptually related to approaches that directly capture physical cell–cell conjugates, including PIC-seq^25^ and more recently cytometry-based interaction mapping frameworks such as Interact-omics^26^. However, these methods typically recover contacts retrospectively (e.g., from dissociated samples or after bulk co-incubation), such that interaction onset and duration are heterogeneous and cannot be experimentally synchronized; moreover, sample handling can disrupt fragile synapses and/or introduce technical aggregates that blur truly native contacts. In contrast, by engineering and preserving physical pairing, Cell-Cell-seq synchronizes the onset of interaction and maintains dyad identity through handling and sorting. We further isolate interaction-driven programs by benchmarking true dyads against pseudo-dyad controls, that define the expected distribution under simple mixing, and then assign emergent transcripts to their cellular origin to enable partner-resolved interpretation of bidirectional responses within each dyad.

Notably, comparison to ligand–receptor predictions highlights that interaction-induced transcription extends well beyond canonical “communication axes.” In a supplemental Venn analysis (Supplementary Figure 11), a subset of emergent Cell-Cell-seq genes overlaps with the union of ligands and receptors prioritized by CellChat from either co-cultured T cells + PC3 cells or pMHC-activated T cells + PC3 cells (on the order of tens of genes), whereas >300 emergent genes are detected exclusively by Cell-Cell-seq and are not represented among curated ligands/receptors in either baseline. This divergence is expected: ligand–receptor inference prioritizes potential signaling based on expression of known ligand/receptor pairs, while Cell-Cell-seq captures the downstream, context-dependent transcriptional programs that arise from contact, synapse formation, and reciprocal feedback.

When benchmarked against standard well-plate co-culture, Cell-Cell-seq captured transient early response genes such as *IER3*, *EGR1*, *EGR3*, and *NR4A3* more consistently, transcripts known to be induced rapidly following TCR engagement and to decay within hours as activation programs progress^27^. Classical descriptions of these immediate-early waves often rely on synchronized stimulation (e.g., plate-bound anti-CD3/anti-TCR antibodies) to impose a defined onset of signaling across a population^28^. In contrast, target-cell co-cultures initiate asynchronously across many pairs, averaging pre-contact, early-contact, and late-contact states and thereby diluting the short-lived transcripts that best report response initiation. While in vivo timing has been inferred using engineered genetically encoded timer reporters^29^, Cell-Cell-seq synchronizes interaction onset directly by pairing cells and constraining proximity, making these fleeting programs observable under target-recognition conditions and enabling a more mechanistic view of immune response initiation.

At the biological level, Cell-Cell-seq revealed a rich repertoire of emergent interaction programs. Notably, we uncovered a coordinated chemokine bloom, a dyad-level signature in which T cell–derived chemokines (*XCL1*, *CCL3*, *CCL4*) were correlated with tumor-derived chemokines (*CXCL1*, *CXCL3*, *CXCL8*), consistent with bidirectional paracrine signaling. In parallel, we identified a negative correlation module between tumor-expressed *TM4SF1*, associated with immune evasion/immunosuppression in several tumor settings, and multiple T cell activation genes including *CD3D*, *CD3E*, and *CD2*. These compartment-resolved, dyad-linked programs and their putative communication modes are summarized schematically in Supplementary Figure 12. These interactions were absent in pseudo-dyad controls and emerged only in real dyads, indicating that they represent true, dyad-linked cross-cell coupling rather than confounding cell state effects. Given the druggability of factors such as DKK1 and TM4SF1, these dyad-linked signatures suggest opportunities to target immune suppression at the level of individual communication events. Prior work has implicated DKK1 in tumor immune evasion largely through remodeling of suppressive myeloid compartments, with additional evidence for direct inhibition of CD8 T cell programs in some settings^30^. For TM4SF1, recent studies in hepatocellular carcinoma connect tumor TM4SF1 to immune evasion and report impaired CD8 T cell function consistent with a more direct tumor–T cell axis, but dyad-resolved transcriptional coupling has remained difficult to demonstrate with conventional assays^31^.

Comparison to *in vivo* T cell trajectories further demonstrates the utility of Cell-Cell-seq. In an adoptive transfer model using the same TCRs profiled here, single-cell RNA-seq of tumor-infiltrating lymphocytes recovered a pseudo-temporal progression from quiescence to early effector, late effector, and terminal ISG-high states. Cell-Cell-seq recapitulates and compresses features of these trajectories into a rapid, experimentally controlled window, capturing within hours chemokine-rich (*CCL3*, *CCL4*) and interferon-stimulated (*IRF1*, *NFKBIA*) modules that in vivo emerge over days of tumor residence. These results suggest that early dyad-level transcriptional programs measured by Cell-Cell-seq can provide proximal correlates of longer-term in vivo states (e.g., persistence versus dysfunction), motivating future studies that explicitly link early dyadic signatures to downstream fate.

Technically, Cell-Cell-seq builds on and significantly expands prior Nanovial-based approaches, which have been used to profile secretory phenotypes^8,32^, cytokine output^33^, and EV production^34^. The platform uniquely integrates high-throughput dyad formation, functional readouts, and dyad single-cell transcriptomics, offering an experimentally tractable and generalizable system for studying immune-tumor, immune-immune, or other intercellular interactions. While other platforms such as DEP-based pairing or microfluidic chips offer control over cell pairing, they often lack scalability, single-cell transcriptomic resolution, modularity for suspension and adherent cells, or compatibility with droplet-based sequencing^35–39^. Cell-Cell-seq directly fills this gap.

Finally, we addressed a core technical limitation of dyad sequencing, the compositional dilution in mixed transcriptomes, using ccRepair. By rebalancing dyad contributions based on anchor genes learned from singlets, ccRepair mitigates systematic skew in apparent cell-type dominance, restores suppressed effector transcripts, and improves quantitative comparisons of activation intensity across dyads. Together, these experimental and computational advances position Cell-Cell-seq as a complementary approach to inference methods: it retains their hypothesis-generating value while providing a direct, paired readout of interaction-induced programs, including large modules that are invisible to ligand–receptor catalogs alone.

Beyond T cell–tumor interactions, the platform is readily extendable to other systems including APC–T cell, stromal–immune, B–T cell, or stem–niche interactions by altering capture ligands on the Nanovials^40^. Integration with other modalities such as CITE-seq^41^, ATAC-seq^42^, or CUT&Tag^43^ will further enrich the platform’s multiomic depth. For example, combining Cell-Cell-seq with surface proteomics would allow tracking of immune checkpoint receptor engagement and downstream transcriptional response within a dyad. Similarly, linking chromatin accessibility profiles to interaction timing could reveal early regulatory events driving fate decisions such as exhaustion or memory formation.

From a translational perspective, Cell-Cell-seq offers a framework for functional screening of cell-cell interaction therapies, such as TCRs, chimeric antigen receptors (CARs), bispecific antibodies, or checkpoint inhibitor antibodies. By linking transcriptional responses and cytokine output to specific interactions, it enables both discovery and prioritization of potent therapeutic candidates in a native cell–cell engagement context with all of the associated heterogeneity of each of the interacting partners. Furthermore, by correlating early dyadic signatures with long-term outcomes (e.g., persistence, killing, or exhaustion), Cell-Cell-seq could uncover predictive biomarkers of functional potency, guiding the rational design of cell-cell interaction therapies.

In summary, Cell-Cell-seq introduces a foundational technology for dissecting cell–cell communication with temporal, functional, and transcriptional resolution. It bridges the gap between single-cell sequencing and spatially contextualized interaction analysis, providing a scalable and versatile approach for understanding how cells recognize, respond to, and influence each other. As the field advances toward personalized cell-cell interaction therapies, Cell-Cell-seq offers a unique and timely tool to chart the molecular choreography of immune recognition, therapeutic response, and intercellular signaling, one dyad at a time.

## 4. Methods

### 4.1. Nanovial Preparation and Cell Loading Optimization

Nanovials were fabricated utilizing previously described protocols^6^. Aqueous gelatin solution was flowed into a three-inlet flow-focusing microfluidics droplet generator where it was mixed with a second aqueous solution consisting of 4-Arm-PEG 5KDa and Lithium phenyl-2,4,6-trimethylbenzoylphosphinate) photoinitiator (LAP, Sigma Aldrich). The mixture was then introduced to an oil-phase composed of Pico-surf (Sphere Fluidics) surfactant in Novec 7500 (3M) at the junction where the droplets were generated. A phase equilibrium resulted in separation of the two aqueous phases within the droplet. The phase-separated droplets were UV-crosslinked downstream the generator and immediately before the collection of the now-crosslinked particles in a tube. Next, the particles were washed to remove unpolymerized phases. The cavity of the resulting Nanovials was then specifically decorated with NHS-conjugated biotin (ApexBio) generating streptavidin binding sites for modular surface functionalization. Finally, the Nanovials were sterilized and stored in an aqueous buffer (washing buffer) consisting of Phosphate-buffered Saline solution (PBS, Gibco), 1% Antibiotic-Antimycotic (Gibco), 0.05% Pluronic F-127 (Sigma Aldrich), 0.05% Bovine Serum Albumin (BSA, Sigma Aldrich) at 4°C until usage. The inner cavity size for the 48µm ± 2µm Nanovials fabricated were 35µm ± 1µm. Nanovials were either utilized unmodified as described above or were functionalized in a two-step process as previously described^6^ to study the secretion profile of the interacting cells. Utilizing the pre-existing biotinylated sites, streptavidin solution was first added to the Nanovials, incubated at room temperature on a slow rotator for 30min and then washed three times with excess washing buffer. In the final step, Nanovials were incubated with biotinylated anti-human granzyme B antibody (R&D systems) or PAP pMHC monomer at room temperature on a slow rotator for 30min and then washed three times with excess washing buffer.

In order to maximize the efficiency of loading single PC3-T cell dyads on Nanovials, a few optimization experiments were performed. These experiments aimed to determine the optimum cell-to-Nanovial ratio for loading each cell sequentially. Nanovials were initially mixed and loaded with PC3 cells at various cell-to-Nanovial ratios (0.5, 1.5, 2, 3, and 4.5) and the resulting samples were analyzed with flow cytometry. The percentage of PC3-loaded Nanovials increased with increasing cell-to-Nanovial ratios, reaching a peak at a ratio of 3 and then plateauing at higher cell-to-Nanovial ratio (Supplementary Figure S1). Next, the efficiency of capturing T cells onto Nanovials pre-seeded with PC3 cells were accessed. A fixed number (0.2 million) of PC3-loaded Nanovials were mixed with varying numbers of T cells to achieve different T cell-to-Nanovial ratios (0.5, 1, 1.5, 2, and 2.5) and the efficiency of each sample was determined utilizing flow cytometry analysis. The percentage of Nanovials loaded with both cell types increased with higher T cell-to-Nanovial ratios up to 1.5 and exhibited no significant improvement at higher ratios of 2, and 2.5 (Supplementary Figure S1). The optimized sequential loading protocol resulted in efficient co-loading of PC3 and T cells onto Nanovials, maintaining stable cell attachment throughout subsequent handling and FACS processing steps. This stability is crucial to preserve accurate analysis and reliable downstream functional assessment of the interacting cell dyads. Therefore, the PC3 cell-to-Nanovial ratio of 3 and the T cell-to-PC3-loaded Nanovials ratio of 1.5 was selected for all future experiments.

Using the optimal conditions described above, cells were loaded into Nanovials sequentially by first mixing the GFP+/RFP+ expressing prostate cancer cells expressing PAP antigen (PC3-A2-PAP) with 48µm Nanovials at a 3:1 cell to Nanovial ratio in a 24-well plate pre-filled with 1.5ml target cell medium (comprised of 10% FBS (Thermo Fisher Scientific) and 5% GlutaMAX supplement (Thermo Fisher Scientific) in Ham’s F-12K base medium (Thermo Fisher Scientific). Next, the well plate was incubated for 2hrs in an incubator at 37°C and 5% CO_2_. At 30min intervals, the plate was removed from the incubator and the cell-Nanovial mixture was re-suspended with a 1ml pipette 10 times. At the 2hr time point, the cell-Nanovial mixture was passed through a 20µm reversible strainer (CellTrics) and washed with an excess amount of washing buffer to remove unbound tumor cells. This resulted in loading the tumor cells into the Nanovial cavities leveraging the pre-existing gelatin layer (which served as the binding motif for the tumor cells). The strained Nanovial suspension was then recovered in another 24-well plate with 1.5ml target cell medium. Next, pre-stained (CellTracker deep red) T cells (expressing cognate TCR) were added to the well in a 1.5:1 cell to Nanovial ratio. The well plate was incubated for 1.5 hours in an incubator at 37°C and 5% CO_2_ with a pipetting re-mix step at the intermediate time point. At the end of the incubation period, the cell-Nanovial mixture was passed through a 20µm reversible strainer (CellTrics) and washed with an excess amount of washing buffer to remove unbound T cells. This resulted in loading the cognate TCR T cells into the Nanovial cavity via their capture by the pre-seeded tumor cells within the Nanovial cavity. Finally, Nanovials pre-seeded with PC3 cells were loaded with non-cognate TCR T cells to test the specificity of the PC3-TCR T binding.

### 4.2 Flow Cytometry Gating of Nanovials with Cell Dyads

PC3-T dyad-loaded Nanovial samples were analyzed to isolate the population of interest and sorted with the Sony SH800S flow sorter. In order to isolate the cell dyad-loaded Nanovials, first the SSC-A vs FSC-H plot was utilized to isolate the Nanovial population. Next, the FITC-A vs FITC-W signals plot was employed to isolate the GFP+ expressing PC3 cell-loaded Nanovials from the empty Nanovial population, and to exclude the undesired population of i) free cells (appearing on the left side of PC3-loaded Nanovials gate) and ii) Nanovial aggregates (appearing on the right side of the PC3-loaded Nanovials gate). Finally, the cell dyad-loaded Nanovial population was isolated via drawing a gate on the APC-A vs APC-W plot. To determine the ability to isolate accurate cell dyads utilizing this gating strategy, the resulting cell dyad-loaded Nanovials were sorted into a 96 well plate and imaged using an inverted fluorescence microscope (Nikon ECLIPSE Ti). Image analysis of the sorted populations was used to confirm the successful identification and isolation of the desired population.

### 4.3. Functional Characterization of T Cell Activation in Dyad-loaded Nanovials

Nanovials conjugated with anti-human granzyme B antibodies were prepared and sequentially loaded with PC3 tumor cells and T cells (TCR156-45 or wild-type) as described earlier. In order to identify the cytokine secretion signal threshold and exclude background signal, a sample composed of Nanovials without cytokine capture antibodies was also prepared and loaded sequentially with both cells. Finally, two positive control samples were prepared in parallel; Nanovials conjugated with PAP pMHC monomer and loaded with TCR156-45 T cells as well as Nanovials conjugated with anti-human CD45 antibody (BioLegend) and loaded with TCR156-45 T cells (for PMA-ionomycin activation later on). After loading the cells, the end of the second straining step described earlier in the cell-loading protocol, each experimental sample was recovered with a 2ml target cell medium in a 6-well plate (the CD45-coated Nanovial sample was recovered with 2ml target cell medium containing 2 µl of Cell Activation Cocktail (BioLegend) composed of PMA, ionomycin, and Brefeldin A). The well plate was then incubated at 37°C and 5% CO_2_ for 12 hours to allow the capture of interaction induced secreted cytokines on the associated Nanovials. Finally, the samples were collected in a 5ml Eppendorf tube, stained with BV421 anti-human granzyme B (BioLegend) and AF647 anti-human CD271 (NGFR) detection antibodies (BioLegend), washed one time with excess washing buffer, and resuspended in the buffer at 500 Nanovials per µl for flow analysis on the Sony SH800S sorter. The double-cell-loaded Nanovials were isolated based on the gating strategy described earlier. The secretion signal threshold was set based on the background signal level from Nanovials without capture antibodies sample.

In order to access the granzyme B secretion profile over time for the PC3-cognate TCR T dyad Nanovials, the secretion experiment described here was repeated for varying incubation periods of 3, 7, and 11 hours time points. The detection antibodies utilized for this assay include BV421 anti-human granzyme B (BioLegend) and APC/Fire750 anti-human CD271 (NGFR) detection antibodies (BioLegend). The samples were also analyzed utilizing the same gating and threshold sample preparation and set up as described for the previous experiment.

### 4.4 Library Preparation using Single-cell-sequencing of Nanovial Cell Dyads

Two Cell-Cell-seq runs were performed using the 10x Chromium GEM-X platform for library preparation of cell dyad mixed transcriptomes. For the first set (run 1), four distinct experimental samples were prepared; i) T cell (TCR156-45) singlets (captured on Nanovials cavity via CD45 coating) ii) PC3 cell singlets (captured through interactions with the gelatin layer in Nanovial cavities) iii) activated T cell (TCR 165-45) singlets (captured and activated via pMHC monomer coating in the Nanovial cavities) and iv) the PC3-T cell (TCR 165-45) interacting dyad sample. The first two samples, (i) and (ii), served as control conditions to obtain the baseline transcriptomics information of the isolated T cells and PC3 cells without interaction. The Nanovials were loaded with PC3-A2-PAP cells (passage 11) and then T cells (TCR156-45, day 9) utilizing the cell loading protocol described earlier. T cells were stained with CellTracker Deep Red (Thermo Fisher Scientific) before loading. After straining of unbound cells, the last step of the cell loading protocol, each sample was transferred to a 6 well plate with 2ml media and stored in an incubator at 37°C and 5% CO_2_ for a controlled interaction time of 3 hours. At the end of the incubation period, the samples were collected in a 5ml tube and washed with excess washing buffer once. The samples were then resuspended in washing buffer at 500 Nanovials per µl for flow analysis and sorting on the Sony SH800S sorter. Each sample was then enriched with FACS for the population of interest; i) Nanovials loaded with single T cells ii) Nanovials loaded with single PC3 cells iii) Nanovials loaded with single pMHC-activated T cells and iv) Nanovials loaded with T cells and PC3 cells.

For the second Cell-Cell-seq experiment (run 2), the same four samples described for run 1 were prepared with the addition of a fifth sample prepared in parallel; (v) a standard co-culture sample composed of T cells and PC3 cells mixed in 1:1 E:T ratio in a 24 well plate in target cell medium. The co-culture was mixed well and placed in an incubator at 37°C and 5% CO_2_ for 3 hours. At the end of the incubation period, each of the five samples were collected in a 5ml tube and washed with excess washing buffer once. All samples were then stained with APC/Fire750 anti-human CD271 (NGFR) detection antibody (BioLegend) to label T cells. This reaction was performed on ice for 30 minutes followed by washing one time with excess washing buffer. The Nanovial samples (i-iv) were resuspended in washing buffer at 500 Nanovials per µl while the standard sample was resuspended at 1000 cells per µl for flow analysis and sorting on the Sony SH800S sorter. Sample (i-iv) were enriched for the sample populations of interests as described for run 1. For the standard co-culture sample (v), an equal number of GFP+ PC3 cells and NGFR+ T cells were sorted into a tube.

At the end of the sorting process for each run, each sample was then processed separately for 10x single-cell 3’GEX library construction (with GEM-X chemistry) following the manufacturer’s protocol for single-cell analysis, where Nanovials were substituted as single cells. The libraries were then sequenced on 0.5 lanes on the NovaSeq X Plus 10B (2×50) for run 1 and on 1 lane of NovaSeq X Plus 10B (2×50) for run 2.

### 4.5 Cell Culture and Engineered TCR-T Cell Production

TCR-engineered cells used for the assay and analysis were cultured and processed as previously described^10^. Commercially available human peripheral blood mononuclear cells (PBMCs) from HLA-A*02:01 healthy donors were obtained from AllCells (Discovery Life Sciences Inc). Cryopreserved PBMCs were activated overnight with CD3/CD28 Dynabeads (Thermo Fisher Scientific) and cultured in T cell medium (TCM) consisting of AIM V medium (Thermo Fisher Scientific), 5% human AB serum (Sigma-Aldrich), 50 U/mL recombinant human IL-2 (PeproTech), 1 ng/mL recombinant human IL-15 (PeproTech), 1× GlutaMAX (Thermo Fisher Scientific), and 50 μM β-mercaptoethanol (Sigma-Aldrich). Dynabeads were added at a ratio of 25 μl per 1 x 10⁶ cells.

Two rounds of spin infection with MSGV retrovirus encoding the TCRs of interest, prepared according to a previously established protocol^9^, were performed on days 1 and 2 at 1,350 x g for 90 minutes at 30 °C. On day 3, PBMCs were washed once with phosphate-buffered saline (PBS) and returned to culture in fresh TCM. Dynabeads were removed on day 5. On day 7, flow cytometry was performed to assess cell phenotype (human CD3 and human CD8), transduction efficiency using truncated nerve growth factor receptor (NGFR), and membrane trafficking of the introduced TCRs (murine TCRβ).

### 4.6 Computational Analysis

#### Data preprocessing and UMAP visualization

We analyzed transcriptomic profiles from four Nanovial sample types: PC3 cells alone, unactivated T cells alone, PC3–T cell interacting dyads, and activated T cells alone. Low-quality cells and lowly expressed genes were removed, so that the retained cells contained at least 500 detected unique molecular identifier (UMI) counts and less than ten percent mitochondrial gene expression, and the retained genes were detected in at least ten cells. PC3–T cell interacting dyads were retained if they contained at least 10% proportional expression of private genes from both cells, indicating a dyad contains one T cell and one PC3 cell.

UMAP visualization was performed using Seurat (version 4.4.3). After the above filtering steps, the data were normalized and scaled, two thousand highly variable genes were selected, and principal component analysis was conducted on these highly variable genes. The top twenty principal components were used to compute the UMAP embedding, which visualized the four cellular populations (differences in sequencing depth were accounted for by the prior normalization step).

#### Gene–gene correlation networks and signature gene selection

Gene–gene correlation networks were constructed with GeneChorus (v0.12.5) using normalized expression values across cells and interacting dyads. For each gene pair, Pearson correlation coefficients were computed across clustered cells in scaled-squared space over progressive rounds, and edges were retained if the squared correlation coefficient exceeded 0.1. GeneChorus modules were summarized as gene sets and annotated by Gene Ontology enrichment of the genes within each module; modules are referred to in the text by their top enriched gene ontology terms. To prioritize modules for visualization, we computed the mean expression of each module’s genes within each Seurat-defined cluster and selected modules showing the strongest cluster-specific enrichment (highest mean module expression relative to other clusters). For downstream visualization and scoring, we defined “signature genes” as the top 25 genes by absolute log2 fold-change from differential expression of PC3 versus resting T cells (Seurat Wilcoxon test), or alternatively the top 25 highest-degree genes within selected correlation modules (as specified in the relevant figure legends).

#### Pseudo-dyad construction and assignment of emergent genes to PC3 or T cells

To distinguish interaction-induced (“emergent”) transcriptional changes from compositional effects of mixing two transcriptomes, we generated pseudo-dyads by in silico combining single-cell profiles and compared them to measured PC3–T cell interacting dyads (Supplementary Figure 7).

*Pseudo-dyad generation.* We constructed two pseudo-dyad sets: resting pseudo-dyads (PC3 + unactivated T cells) and activated pseudo-dyads (PC3 + activated T cells). For each set, we first converted raw UMI counts to relative expression by dividing each gene count by the total UMI count per cell. We then repeatedly sampled equal numbers of PC3 and T cells (n = 895 each; 895 corresponds to the smallest group size among PC3 singlets, unactivated T singlets, activated T singlets, and interacting dyads) and formed pseudo-dyads by averaging paired profiles:

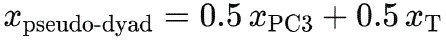

Each resampling produced a dataset of 895 pseudo-dyads; we generated 50 independent pseudo-dyad datasets per condition (resting and activated). In parallel, we generated 50 matched interacting-dyad datasets by randomly subsampling 895 real dyads per dataset to maintain comparable sample sizes across comparisons.

*Differential expression to identify emergent genes.* For each of the 50 resting pseudo-dyad datasets, we concatenated the pseudo-dyads with one size-matched interacting-dyad dataset and applied Seurat normalization (NormalizeData, scale.factor = 10,000) followed by log transformation (log1p). We then performed differential expression (DE) testing between interacting dyads vs resting pseudo-dyads using Seurat’s Wilcoxon rank-sum test (FindMarkers, test.use = “wilcox”) with Benjamini–Hochberg correction. Genes with adjusted p < 0.05 were recorded for each comparison. Resting emergent genes were defined as those significant in all 50 resting comparisons, yielding a conservative set of interaction-associated genes robust to subsampling.

*Filtering to attribute emergent genes to PC3 versus T cells.* Genes identified from the resting comparison can reflect either (i) T cell activation programs or (ii) PC3 responses to T cell engagement. To help separate these sources, we performed an additional DE comparison between interacting dyads vs activated pseudo-dyads (both containing activated T cells). Because T cells in activated pseudo-dyads were stimulated by pMHC and may not match the activation state of T cells in interacting dyads, this comparison can introduce T-cell-state artifacts. We therefore restricted the resulting DE genes to those plausibly attributable to PC3 by retaining only genes with measurable baseline expression in PC3 cells. Specifically, we removed genes with minimal PC3 expression by testing whether normalized expression in PC3 singlets exceeded 0.05 (one-sided t-test with Benjamini–Hochberg correction) and retaining only genes passing this PC3-expression filter (adjusted p < 0.05).

Finally, we assigned resting emergent genes to cell type as follows: genes present in both (i) the conservative resting emergent set and (ii) the PC3-filtered DE list from interacting vs activated pseudo-dyads were classified as PC3-associated emergent genes. The remaining resting emergent genes were classified as T-cell-associated emergent genes.

#### ccRepair normalization for visualization and correlation analyses

Mixed dyad transcriptomes contain RNA from both cells, which can dilute compartment-specific expression when the two cells contribute unequal transcript abundance. To reduce this compositional bias for visualization and gene–gene correlation analyses, we applied ccRepair, a scaling procedure that rebalances cell-type–specific (“compartment”) genes within each dyad using anchor genes derived from singlet baselines. To account for the dilution effect of mixed transcriptomes in dyads, approximately 100 consistent private compartment genes were selected for pMHC-NV T cells and PC3s (log 2 FC DE > 1.5, normalized expression >1 in target compartment and <0.2 in reciprocal compartment) to serve as anchor genes. These genes were used to calculate a target mean anchor expression in unmixed pMHC-NV cells and PC3 cells, and a scaling factor for correction in interacting dyads. In interacting dyads, the ratio between the average single target anchor gene expression and the average dyad anchor gene value was used as a per-cell multiplier to adjust the diluted gene expression. Only compartment genes were adjusted, and only by their target ratio (PC3 genes with PC3 ratio, T cell genes with pMHC-NV T cell ratio). ccRepair does not affect genes not assigned to a compartment. Adjusted gene expression for interacting dyad cells only were merged back into a normalized count matrix with singlets and unaffected genes.

#### Heatmap

**Figure 4B**. The heatmap was generated from normalized expression of the corresponding genes across six groups: (1) unactivated T cells, (2) PC3 cells, (3) activated T cells, (4) a randomly selected resting pseudo-dyad dataset, (5) a randomly selected activated pseudo-dyad dataset, and (6) PC3–T cell interacting dyads.

**Figure 5D**. The four modules were defined by the gene sets in Figure 3C. Module scores were computed for each interacting dyad using Seurat’s *AddModuleScore()* with the module gene sets as input features. Pearson correlations were then computed between all pairs of module scores and shown as a correlation heatmap.

**Figure 5E**. The heatmap shows Pearson correlations between a subset of T cell private genes and PC3 cell private genes based on their normalized expression in interacting dyads. T cell private genes were defined as T cell emergent genes with no detectable expression in PC3-only cells, and PC3 cell private genes were defined as PC3 cell emergent genes with no detectable expression in T cell-only cells.

The subset of private genes used in Figure 5E was selected as follows:

1. Compute the gene-gene correlation matrix for all private genes using their normalized expression in interacting dyads.
2. Compute the 99.95th and 0.05th quantiles of off-diagonal correlations (upper triangle of the gene-gene correlation matrix) and select gene pairs with correlations above or below these thresholds.
3. Concatenate the relative expression matrices from the 50 resting pseudo-dyad datasets, apply Seurat’s standard normalization, and compute the gene-gene correlation matrix for all private genes in resting pseudo-dyads.
4. Compute the difference between the interacting-dyad correlation matrix (step 1) and the resting pseudo-dyad correlation matrix (step 3).
5. Among the genes involved in the selected pairs from step 2, retain those whose correlation differences (off-diagonal entries) exceeded the 99.5th quantile of off-diagonal entries in the correlation-difference matrix from step 4.

#### Violin plots

**Figure 4C**. The violin plots in the first column were generated from the corresponding genes’ normalized expression across five groups: (1) unactivated T cells, (2) PC3 cells, (3) activated T cells, (4) resting pseudo-dyads (50 datasets), and (5) PC3–T cell interacting dyads. The violin plots in the second column were generated analogously using: (1) unactivated T cells, (2) PC3 cells, (3) activated T cells, (4) activated pseudo-dyads (50 datasets), and (5) PC3–T cell interacting dyads.

**Figures 5F–G**. Interacting dyads were stratified into “CD3E high” and “CD3E low” groups using the ninety-fifth percentile of CD3E normalized expression within interacting dyads as the cutoff: dyads above the cutoff were labeled “CD3E high,” and dyads at or below the cutoff were labeled “CD3E low.” Figure 5F shows TM4SF1 normalized expression in interacting dyads for the two groups. Figure 5G shows TM4SF1 normalized expression in the concatenated resting pseudo-dyad datasets for the two groups.

#### Volcano plots

**Figures 4D–F**. Figures 4D–F display volcano plots for (1) all emergent genes, (2) T cell emergent genes, and (3) PC3 cell emergent genes. For Figures 4D and 4E, the adjusted p-values and log fold changes were computed as the averages across 50 DE tests comparing resting pseudo-dyads with interacting dyads. For Figure 4F, the adjusted p-values and log fold changes were computed as the averages across 50 DE tests comparing activated pseudo-dyads with interacting dyads. In each plot, we labeled (1) the ten upregulated and ten downregulated emergent genes with the smallest adjusted p-values and (2) the five upregulated and five downregulated emergent genes with the largest absolute log fold changes.

#### Scatter plots

**Figure 4G**. We ran Seurat’s default DE test comparing T cells co-cultured with PC3 cells in well plates with unactivated T cells. From the emergent genes identified as described in the “Pseudo-dyad construction and cell-type-specific emergent gene identification” section, we selected the 100 most significantly upregulated and 100 most significantly downregulated emergent genes based on adjusted p-values. We then plotted these genes, together with additional known immediate early genes, using (1) their mean adjusted p-values across the 50 resting pseudo-dyads versus interacting-dyad DE tests and (2) their adjusted p-values from the well-plate co-culture DE test. Emergent genes that were not significant in the co-culture DE test were labeled.

**Figure 5I**. We computed gene-gene correlation matrices for interacting dyads and resting pseudo-dyads as described in the “Heatmap” section and then calculated their difference. Each point in the scatter plot represents a gene pair, showing its correlation in interacting dyads and in resting pseudo-dyads. Gene pairs were formed from T cell private genes and PC3 cell private genes and grouped into three categories: (1) pairs consisting of two T cell private genes, (2) pairs consisting of one T cell private gene and one PC3 cell private gene, and (3) pairs consisting of two PC3 cell private genes. We labeled the top five gene pairs with the largest correlation differences within each of the three categories.

## 5. Acknowledgements

This project has been made possible in part by grant 2023-332386 from the Chan Zuckerberg Initiative Donor Advised Fund (CZI DAF), an advised fund of the Silicon Valley Community Foundation. This work was delivered as part of the MATCHMAKERS team, supported by the Cancer Grand Challenges partnership financed by CRUK (CGCATF-2023/100006) and the National Cancer Institute (OT2CA297242) (K.C.G.), Parker Institute for Cancer Immunotherapy (K.C.G.), National Institutes of Health 5 R01 AI103867 08 (K.C.G.), and Weill Stanford/UCSF Cancer Fund (K.C.G). Parker Institute for Cancer Immunotherapy 20163828, 20221408 (O.N.W.), Parker Institute for Cancer Immunotherapy C-03920 (J.K.L.), National Cancer Institute, UCLA SPORE in Prostate Cancer P50 CA092131 (O.N.W., J.K.L.). Department of Defense W81XWH-20-1-0119 (J.K.L.). UCLA Immunology Advisory Committee Immunology Research Stimulus Award (J.K.L. and D.D.). Deutsche Forschungsgemeinschaft (DFG), Walter Benjamin Fellowship, CH 2718/1-1 (X.C.). UCLA JCCC postdoctoral fellowship (Z.M.). Prostate Cancer Foundation Young Investigator Award 25YOUN17 (Z.M.). Cancer Research Institute Irvington Postdoctoral Fellowship CRI15364 (Z.M.). Partial computational analyses were supported by NSF Grant DGE-2034835 to Q.W. and NIH/NIGMS R35GM140888 to J.J.L. This material is based upon work supported by the National Science Foundation Graduate Research Fellowship Program under Grant No.DGE-2034835. Any opinions, findings, and conclusions or recommendations expressed in this material are those of the author(s) and do not necessarily reflect the views of the National Science Foundation. Sorting experiments were performed in the UCLA Jonsson Comprehensive Cancer Center (JCCC) Flow Cytometry Shared Resource that is supported by the National Institutes of Health award P30CA016042 and by the JCCC and the David Geffen School of Medicine at UCLA. We thank the Broad Stem Cell Research Center (BSCRC) for imaging and UCLA Technology Center for Genomics and Bioinformatics (TCGB) facility for assisting with sequencing. We thank Partillion Bioscience for providing Nanovials in multiple sizes and two distinct formulations, which enabled our compatibility testing with the 10x Genomics GEM-X chip. Selected schematic figures were created using BioRender.

## Supplementary Figures and Figure Captions

**Supplementary figure 1.**
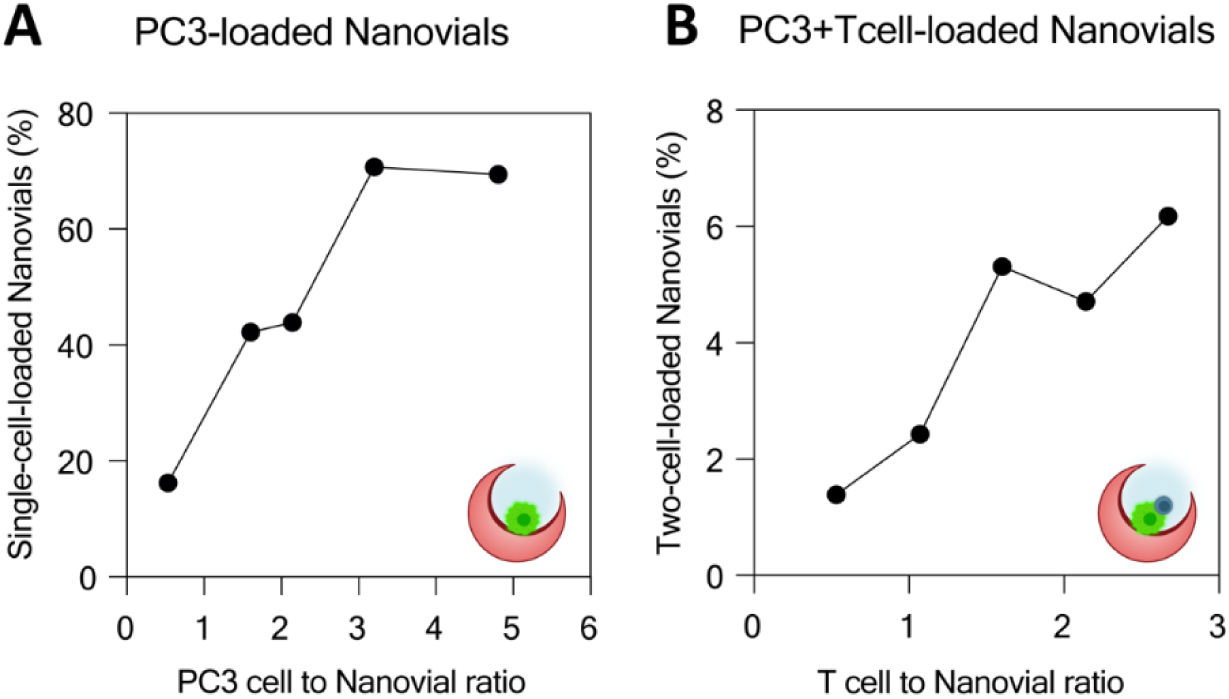
Cell loading optimization of dyad Nanovials. (**A**) Flow cytometry–based quantification of the percentage of Nanovials containing PC3 cells following loading at different PC3 cell–to–Nanovial ratios (0.5, 1.5, 2, 3, and 4.5). (**B**) Flow cytometry–based quantification of the percentage of Nanovials containing PC3-T cell dyads following sequential loading with T cells at different T cell–to–PC3-loaded Nanovial ratios (0.5, 1, 1.5, 2, and 2.5).

**Supplementary Figure 2.**
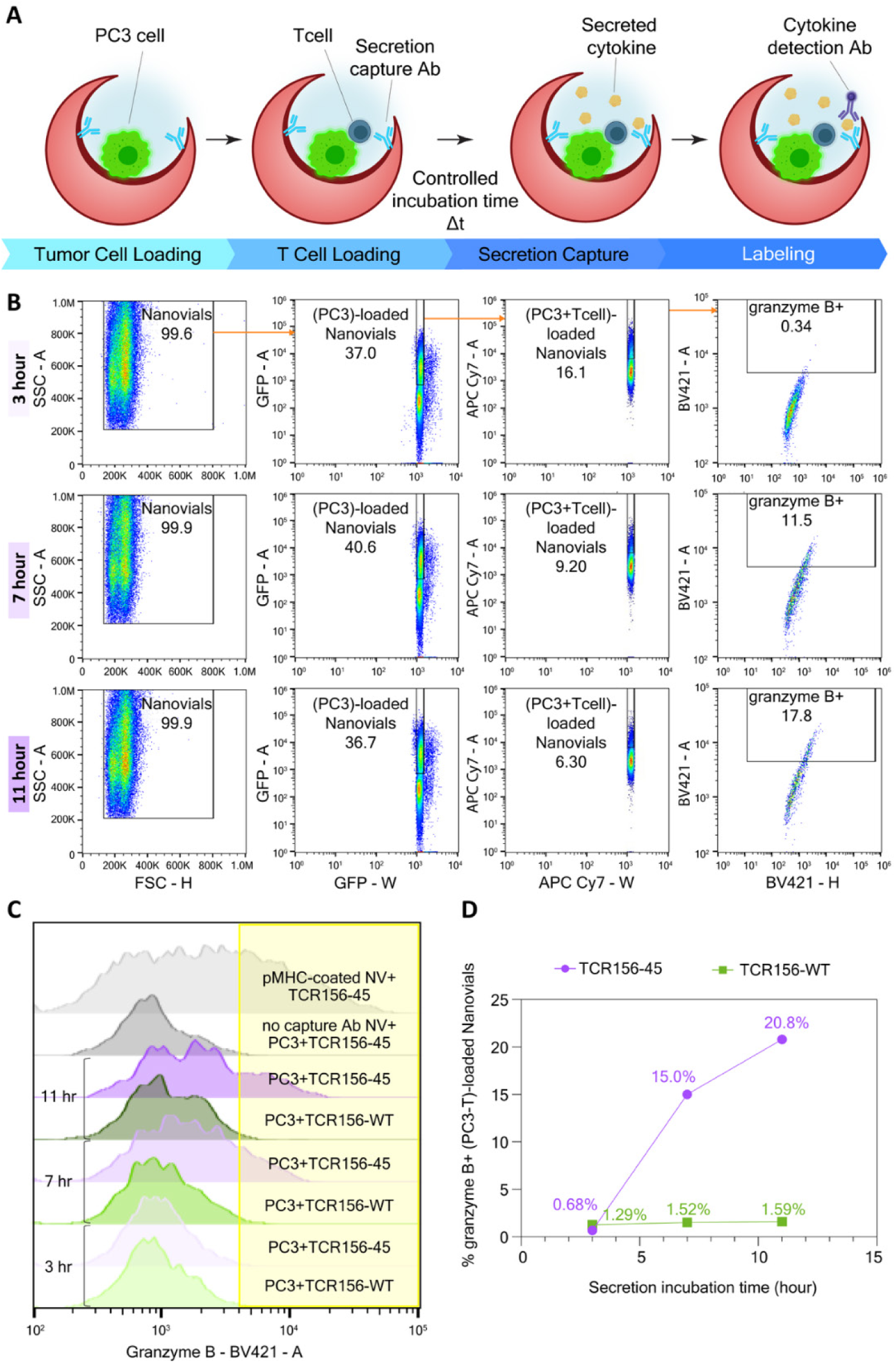
Dynamics of Nanovial granzyme B secretion assay for T cell – tumor cell dyads. **(A)** Workflow schematic. Individual GFP+ PC3-A2-PAP tumor cells are first adhered into respective gelatin-coated Nanovial cavities, followed by capture of T cells (TCR 156-45 or TCR156-WT) through the pre-seeded tumor cells. Secreted granzyme B is captured through capture antibodies on Nanovials during a controlled incubation interval (Δt). Finally, the secreted granzyme B is detected with a fluorescent secondary antibody for flow cytometry. **(B)** Representative flow plots and gating strategy to isolate dyads at three incubation times (3, 7, 11 hours) for (PC3-TCR156-45)-loaded Nanovials. From left to right: identification of all Nanovials (FSC/SSC), enrichment of PC3-loaded Nanovials (GFP+), and detection of PC3-T cell dyads (T-cell marker, APC-Cy7), and granzyme B signal on co-loaded Nanovials. Percentages in gates indicate the fraction of events in each population. **(C)** Overlaid flow histograms of granzyme B signal across time and experimental conditions: (i) no capture antibody (control for setting the signal threshold), (ii) pMHC-coated Nanovials+TCR156-45 (artificial antigen presentation, positive control), (iii) PC3+TCR156-WT (weak TCR), (iv) PC3+TCR156-45 (potent TCR) across 3, 7, 11 hour secretion incubation times. **(D)** Quantification of granzyme B signal among PC3-T cell dyads in Nanovials over time. Plots of the percentages of events with granzyme B above the signal threshold (yellow region) in part (C) are shown for corresponding 3, 7, and 11 hour incubation times.

**Supplementary Figure 3.**
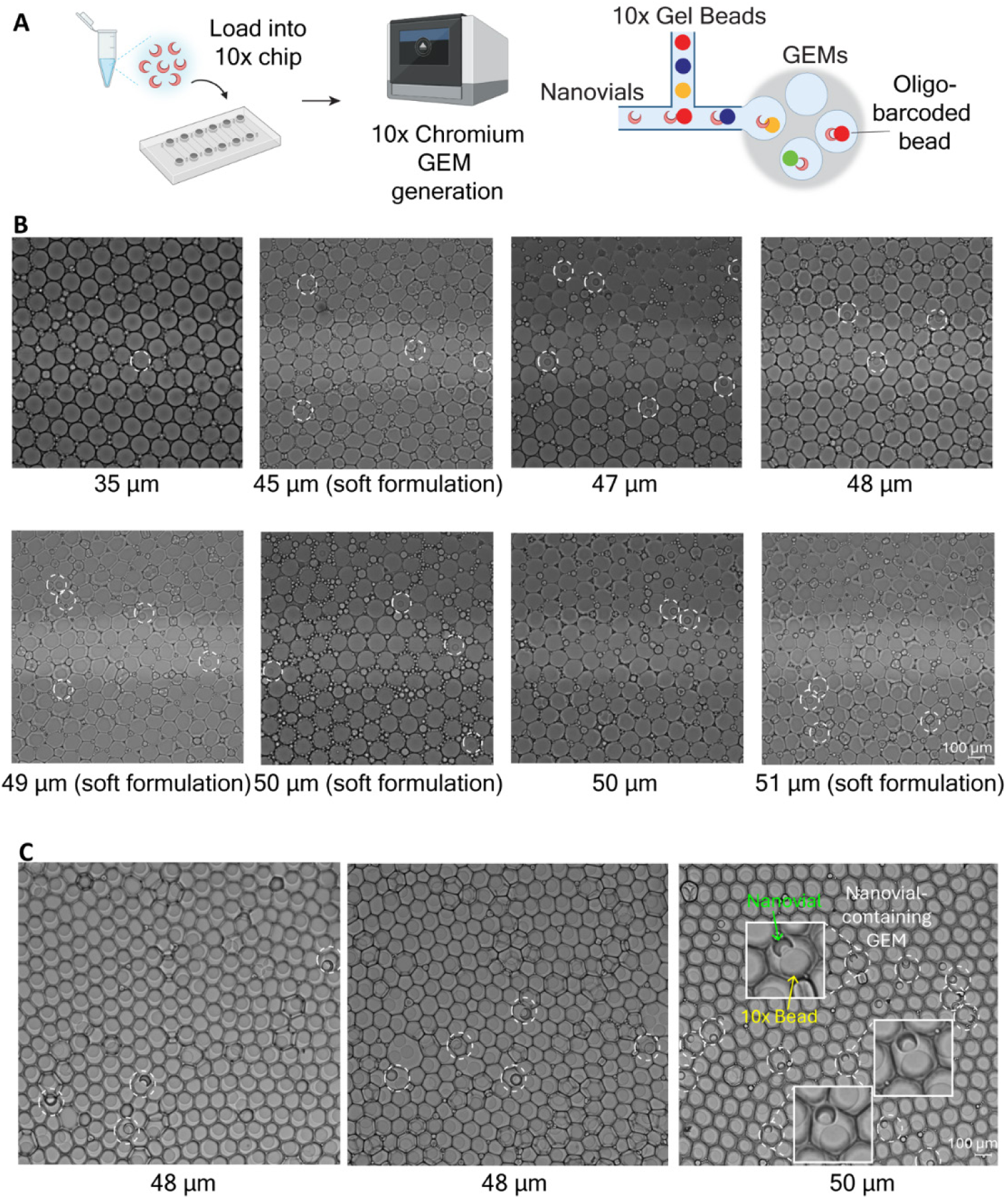
Compatibility of Nanovials with droplet-based single-cell RNA sequencing. **(A)** Schematic illustrating loading of Nanovials into the 10x Genomics Chromium GEM-X chip, insertion of the chip into the Chromium Controller, and formation of gel beads–in–emulsion (GEMs) droplets. The inset shows a schematic zoom of a GEM containing a Nanovial and a barcoded gel bead. **(B)** Representative microscopy images of Nanovials encapsulated within GEMs for Nanovial mean diameters ranging from 35 μm to 51 μm (outer diameter CV of 2.8%–7.1%). Soft formulation refers to PEG formulations with reduced cross-linking density that lead to a less stiff structure. **(C)** Representative images of GEMs containing Nanovials and test gel beads that do not dissolve during emulsification, illustrating co-encapsulation within droplets. Insets show magnified views of individual GEMs highlighting Nanovial–gel bead co-loading.

**Supplementary Figure 4.**
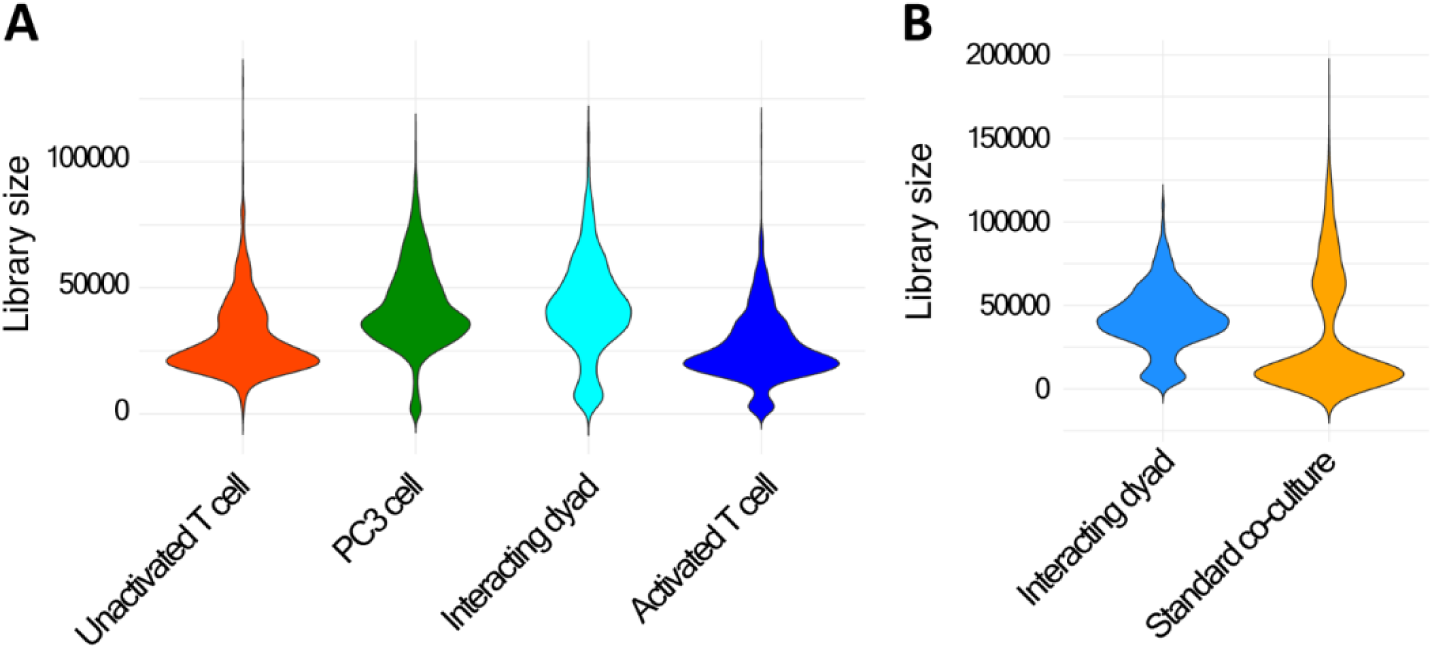
Library size distributions from Cell-Cell-seq and single-cell RNA sequencing experiments. **(A)** Violin plots showing the distribution of transcriptome library sizes for unactivated T cells, PC3 cells alone, interacting PC3–T cell dyads, and T cells activated on pMHC-coated Nanovials, obtained from a Cell-Cell-seq experiment. **(B)** Violin plots showing the distribution of transcriptome library sizes comparing interacting PC3-T cell dyads’ libraries with libraries generated from conventional PC3-T cell co-culture (standard co-culture) RNA sequencing experiments.

**Supplementary Figure 5.**
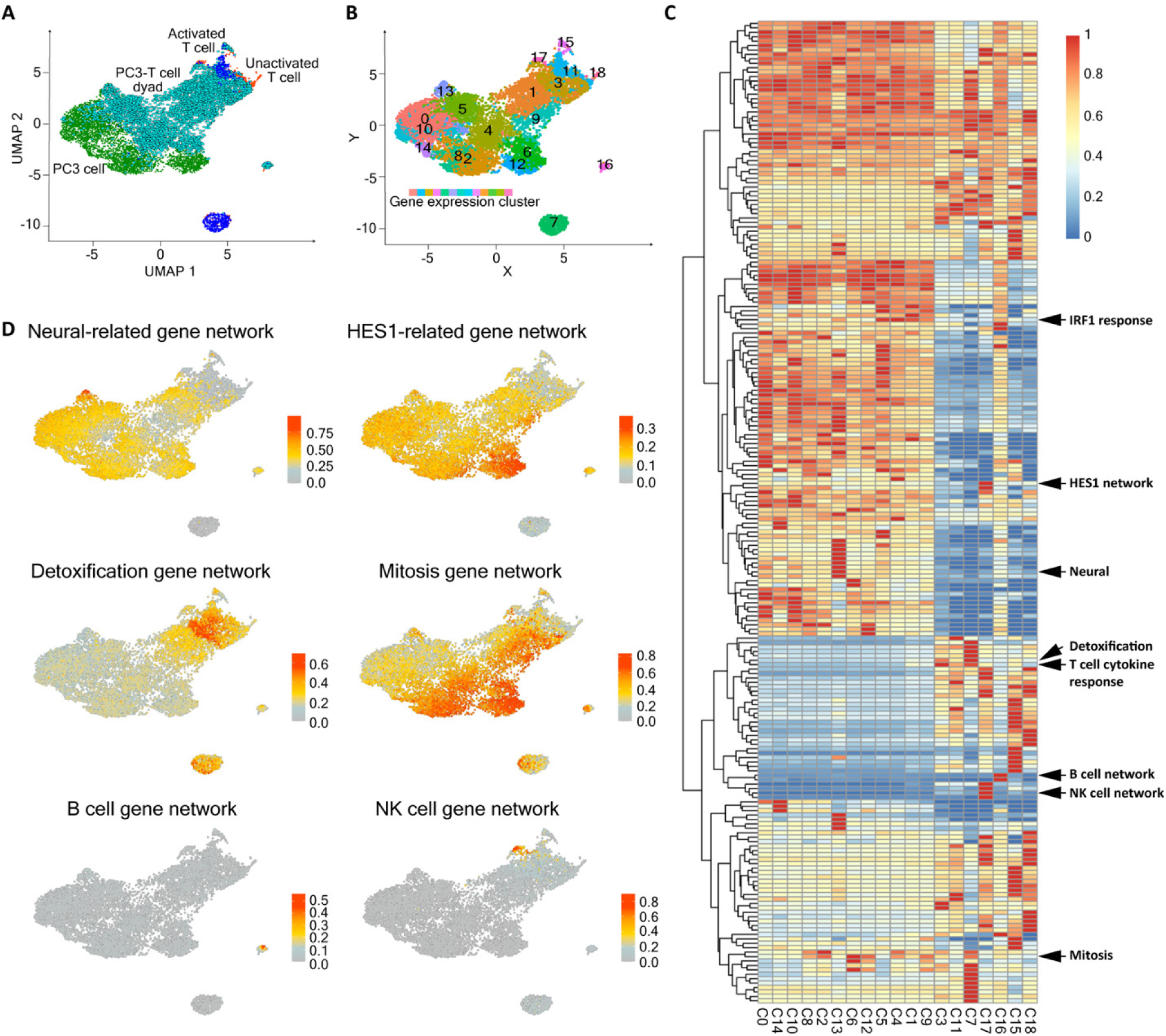
Transcriptional signature enrichment across individual cells and interacting dyad clusters. **(A-B)** UMAP embedding of all Nanovial events of run 1 (left) and corresponding cluster labels (right). **(C)** Heatmap showing transcriptional gene signatures across clusters identified from UMAP embedding. Rows correspond to gene signatures and columns correspond to UMAP-defined clusters. Right-margin annotations highlight the signature families emphasized in the text (figure 3C); IRF1 response, HES1 network, neural-related program, detoxification and T cell cytokine response modules, B cell and NK cell programs, and a mitotic module. **(D)** UMAP projections of interacting dyads with selected transcriptional signature scores overlaid, illustrating the distribution of signature enrichment across individual cells and dyads.

**Supplementary Figure 6.**
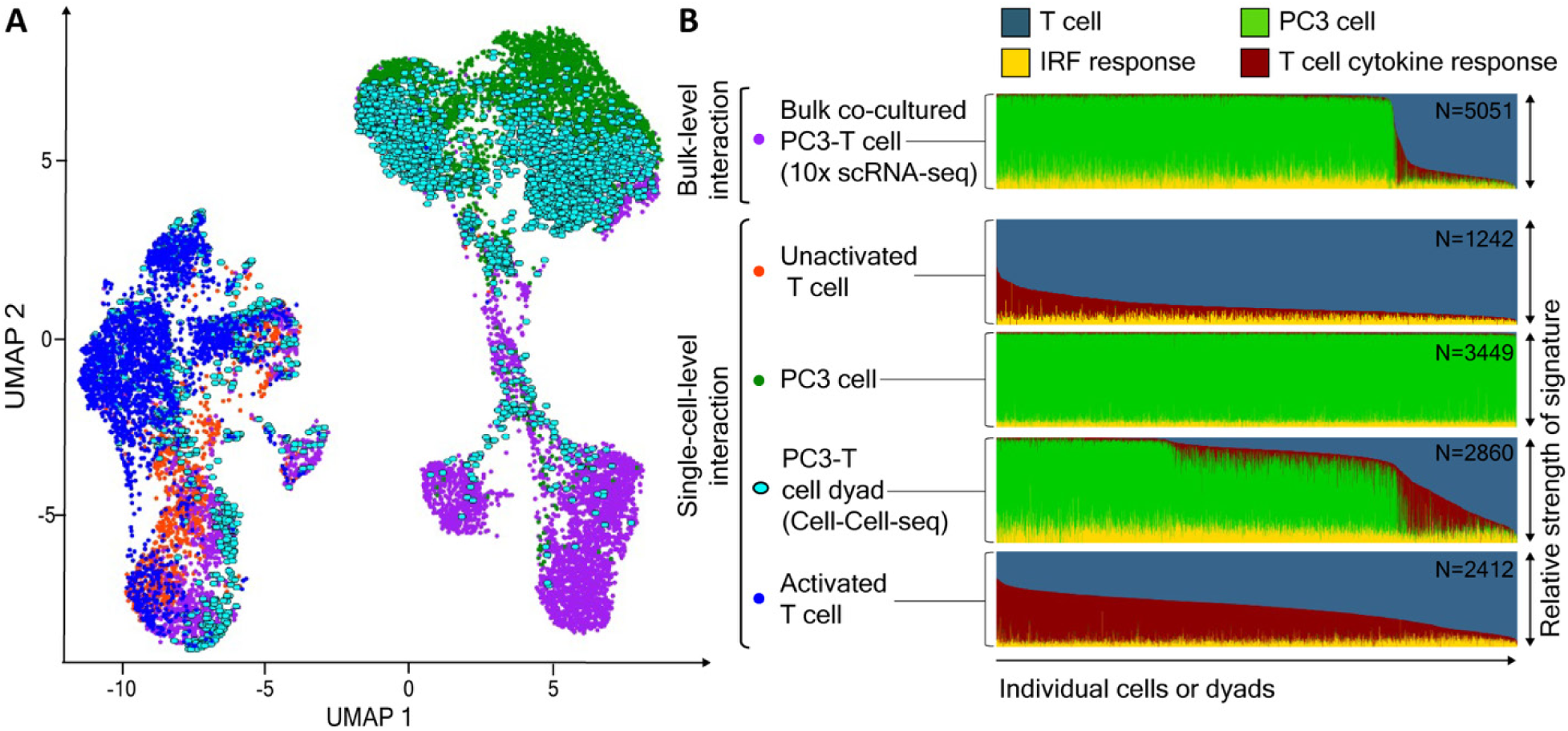
Replicate Cell-Cell-seq experiment and comparison to bulk co-culture. **(A)** UMAP embedding from an independent Cell-Cell-seq experiment (run 2) showing transcriptomes from unactivated T cells, T cells activated on pMHC-coated Nanovials, PC3 cells alone, interacting tumor–T cell dyads, and cells obtained from bulk co-culture of T cells and PC3 cells without controlled pairing or interaction timing, followed by single-cell RNA sequencing. **(B)** Ratiometric representation of transcriptional state scores for individual cells or dyads across the same conditions shown in (A). Each column corresponds to a single cell or dyad, organized along the x axis by similarity in transcriptional state, with conditions stacked along the y axis.

**Supplementary Figure 7.**
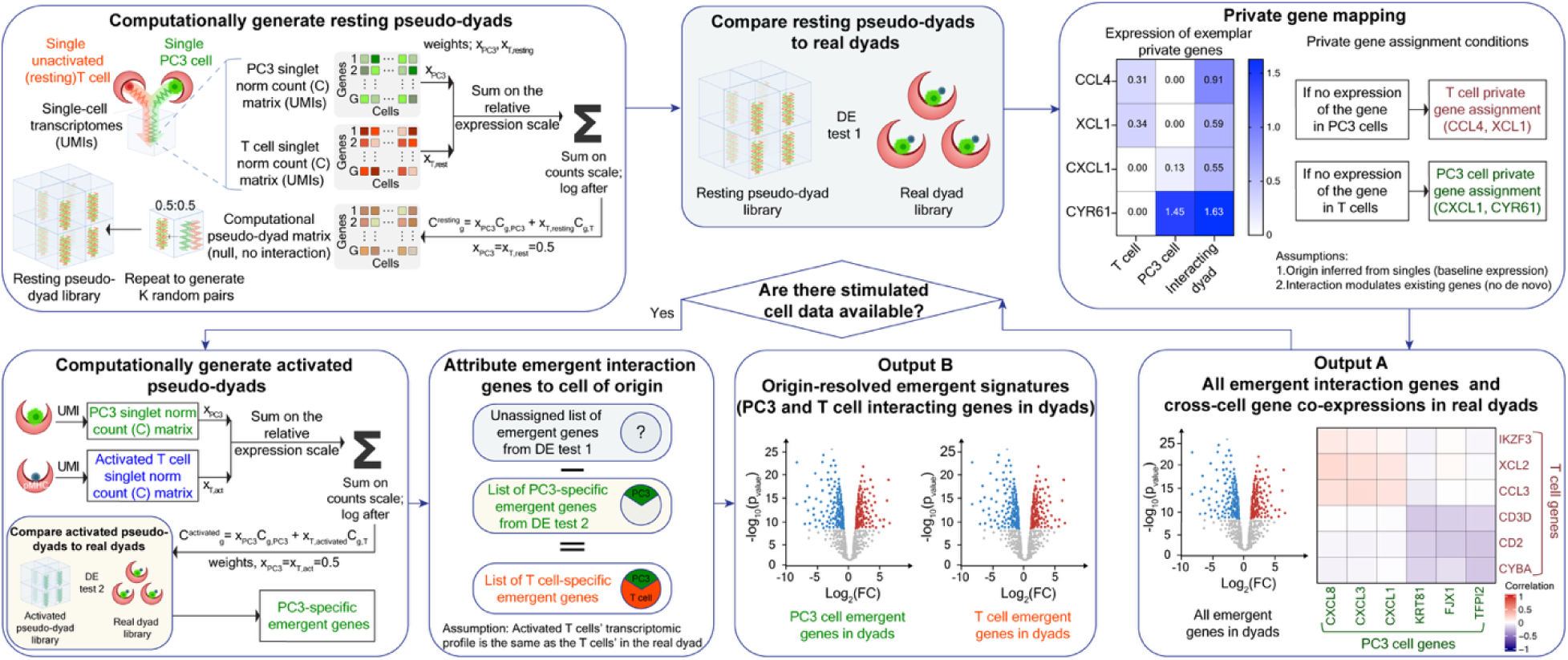
Pseudo-mixing framework. **Computationally generate resting pseudo-dyads.** Single-cell UMI count, C, matrices from PC3 tumor cells and unactivated T cells are paired *in silico*. For each random pair (repeated K times), UMI counts are converted to relative expression levels and summed on the relative expression scale to generate a “null, no-interaction” library. **Compare resting pseudo-dyads to real dyads**. Real dyads are contrasted with the resting pseudo-dyads by Differential Expression test (DE test 1) to identify “emergent” genes. **Private gene mapping.** If stimulated cell (activated T cell) singlet data are not available, emergent genes are provisionally assigned to a cell of origin by private-gene mapping. Genes absent in PC3 singlets are called T cell private; genes absent in T cell singlets are called PC3 private, under the assumptions that (i) origin can be inferred from baseline singlets and (ii) interaction modulates existing programs rather than creating *de novo* expression. These assignments enable cross-cell gene–gene correlation analysis in real dyads in **Output A** (example heatmap) as well as volcano plots without assignment or private gene assignment. **Computationally generate activated pseudo-dyads (optional).** When activated T cell singlets data are available, a second pseudo-dyad library can be built by composing PC3 singlets with activated T cell profiles (depth-matched as in A). Differential expression between real dyads and these activated pseudo-dyads (DE test 2) yields PC3-specific emergent genes. **Attribute emergent interaction genes to cell-of-origin.** The PC3-specific emergent set from DE test 2 is subtracted from the total emergent set from DE test 1; the remainder is assigned as T cell-specific emergent genes, yielding origin-resolved interaction programs which can be displayed as volcano plots as **Output B**.

**Supplementary Figure 8.**
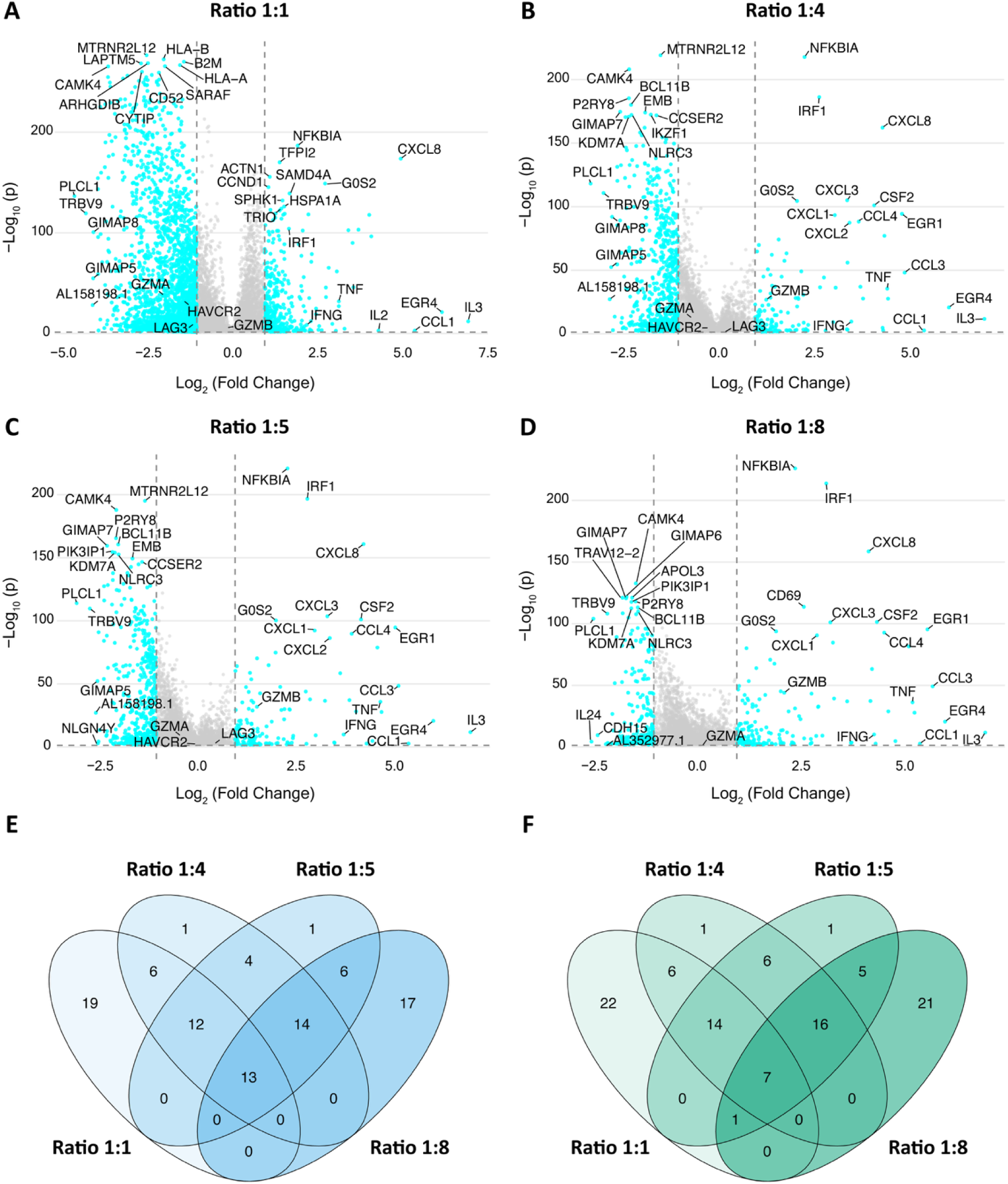
Effect of pseudo-dyad mixing weights on identification of interaction-associated genes. Volcano plots showing differential gene expression between true interacting dyads and computationally generated pseudo-dyads constructed using different transcript mixing ratios. Pseudo-dyads were generated by combining independently sequenced T cell and PC3 cell transcriptomes using the following ratios: **(A)** 1:1, **(B)** 1:4, **(C)** 1:6, and **(D)** 1:8. For each condition, genes are plotted by log₂ fold change versus −log₁₀ adjusted p-value. The top 10 upregulated and top 10 downregulated genes (ranked by adjusted p-values), and the top 5 upregulated and top 5 downregulated genes (ranked by log fold change) are labeled in each panel. **(E–F)** Venn diagrams showing the overlap of the top 50 interaction-associated emergent genes identified using pseudo-dyads generated with each transcript mixing ratio (1:1, 1:4, 1:5, 1:8), separated by inferred cellular origin: (E) T cell–assigned genes (blue) and (F) PC3-assigned genes (green). Numbers indicate the number of genes unique to or shared among ratios.

**Supplementary Figure 9.**
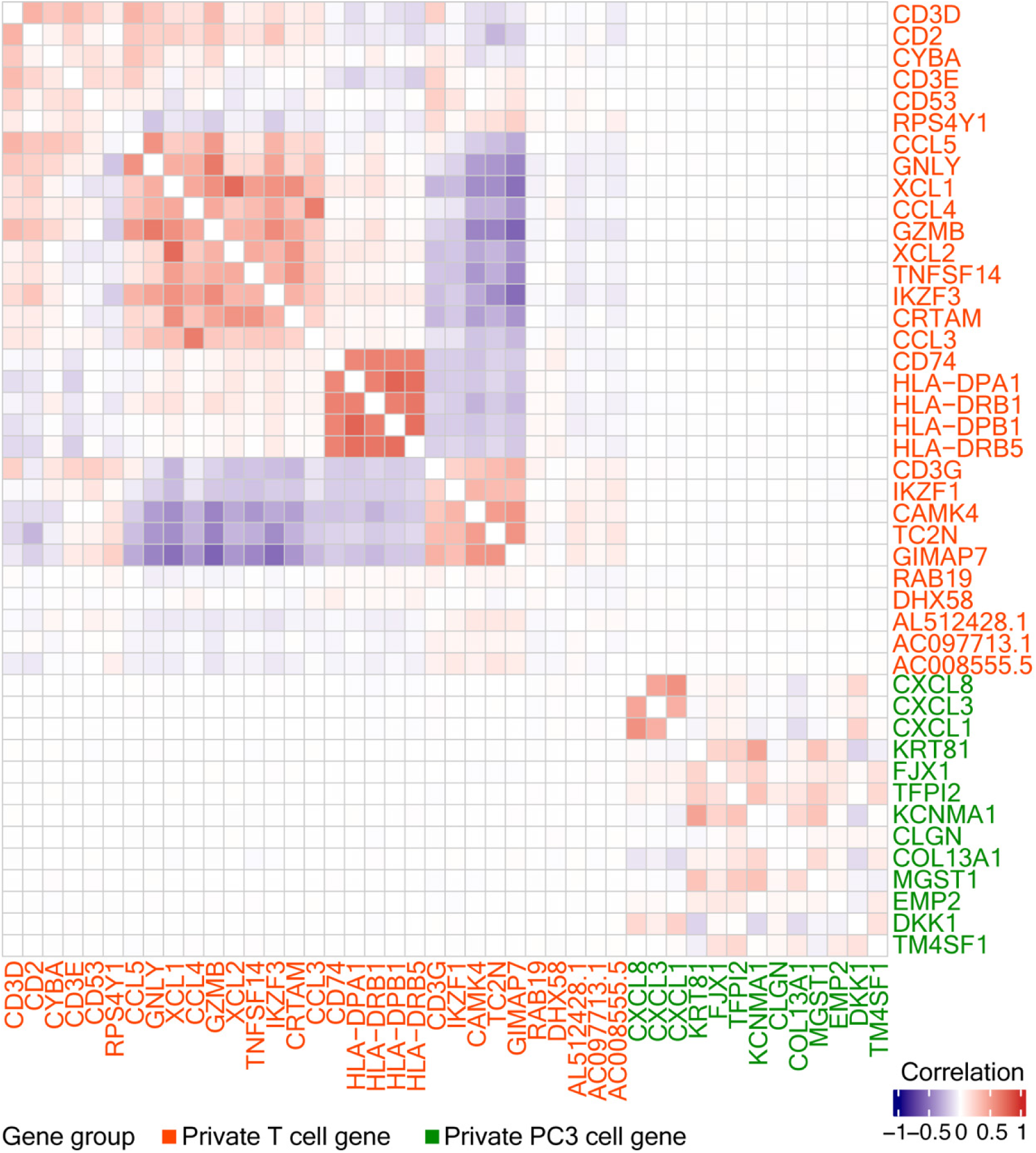
Heatmap showing pairwise correlation coefficients between genes measured across computationally generated pseudo-dyads. The same genes and ordering are used as in Figure 5E, enabling direct visual comparison between correlation patterns observed in true interacting dyads and those arising from pseudo-mixing of independently sequenced T cells and PC3 cells. Correlation coefficients are calculated across all pseudo-dyads and displayed according to the indicated color scale.

**Supplementary Figure 10.**
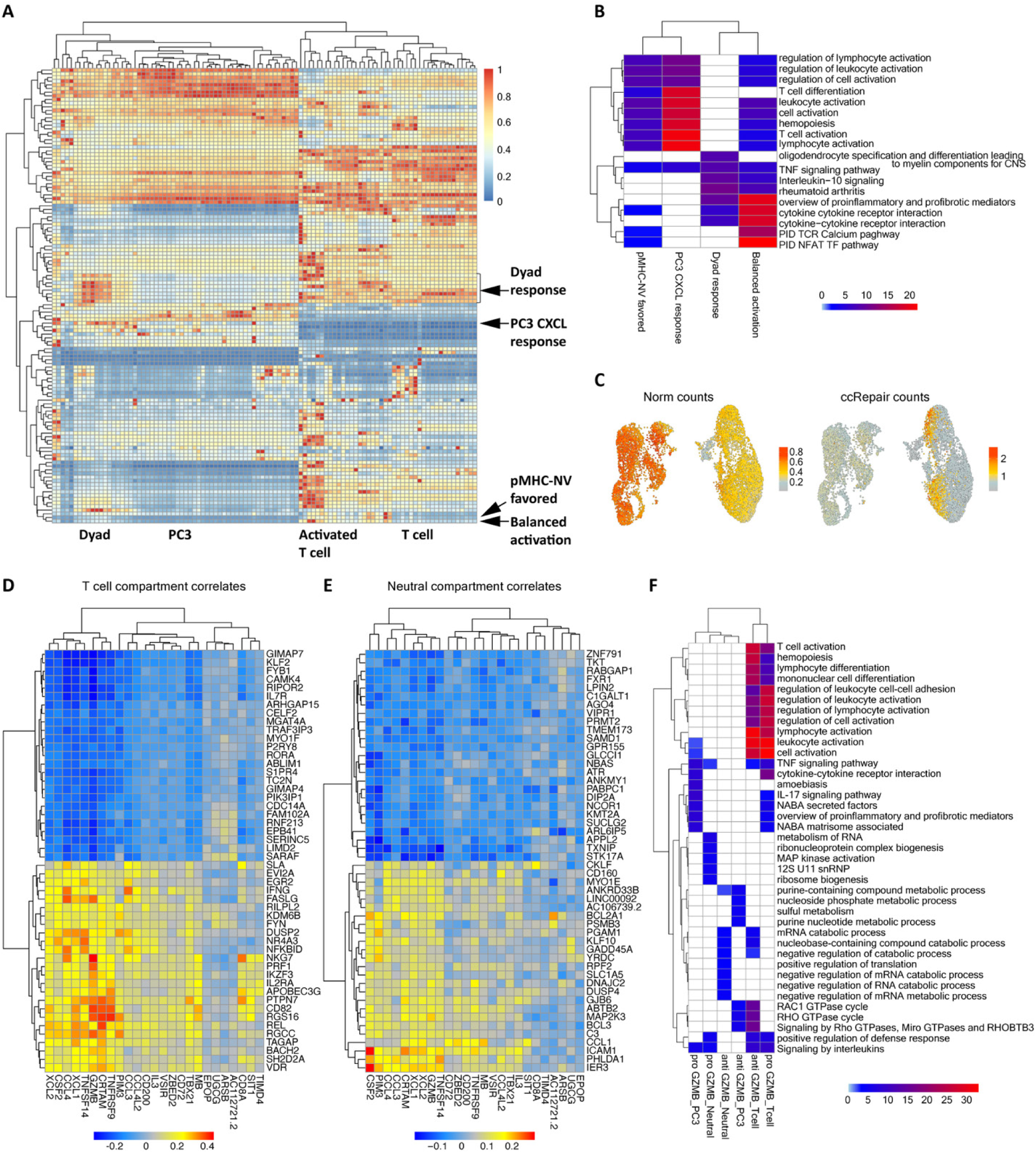
ccRepair sharpens dyad-specific programs and links them to T cell and tumor compartments. **(A)** Gene program heat map (rows represent gene programs, columns represent single cell or dyad groups) after ccRepair. Four modules shown in Figure 6 are identified: dyad response, PC3 CXCL response, pMHC-NV-favored, and balanced activation modules. **(B)** Gene-ontology/pathway enrichment for the four modules in (A). **(C)** UMAPs for the dyad response module (left, uncorrected; right, after ccRepair). **(D-E)** Gene by gene correlation analysis. X-axis genes represent the T cell cytokine response module. Heat maps show the top positively/negatively correlated genes (y-axis) across individual dyads for the T cell compartment (D) and non-T cell or PC3 cell specific compartment genes (neutral compartment correlates) (E). **(F)** Gene ontology/pathway enrichment of the correlation-derived gene sets from (D-E) where GZMB is used as an anchor for positive (pro) and (anti)-correlates.

**Supplementary Figure 11.**
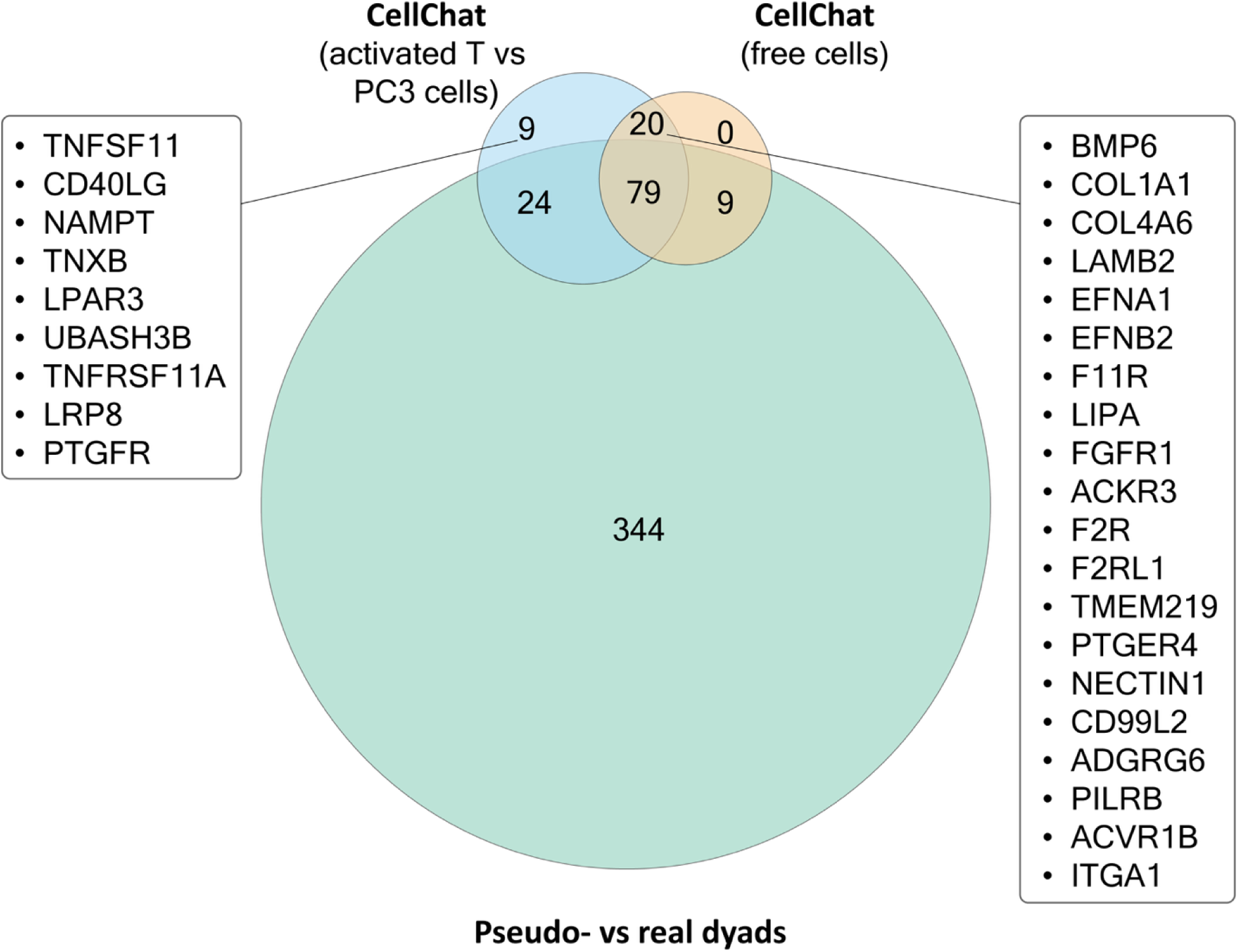
Comparisons between CellChat-inferred interaction genes and Cell-Cell-seq emergent genes. Venn diagram compares the sets of genes highlighted as interaction-relevant by three approaches: CellChat run on activated T cells vs PC3 cells (blue), CellChat run on the free-cell (standard co-culture) populations (orange), and the Cell-Cell-seq pseudo-mixing pipeline (green, emergent genes from real vs pseudo-dyads). Numbers denote the count of ligand / receptor genes in the CellChat library for each intersection.

**Supplementary Figure 12.**
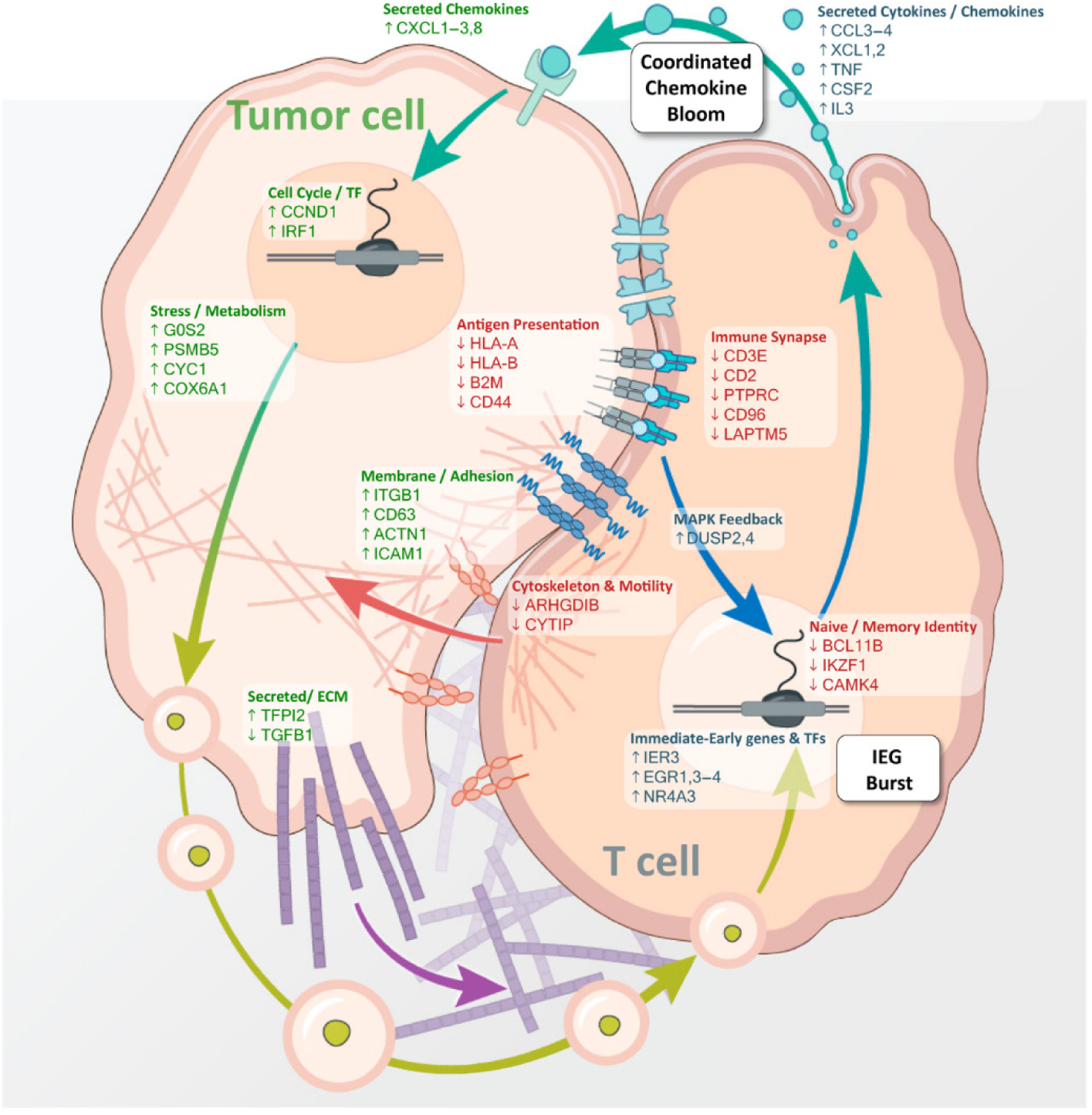
Spatial schematic of compartment-resolved transcriptional programs in interacting tumor–T cell dyads following 4.5 hours of interaction time. Representative interaction-associated genes identified in Cell-Cell-seq are mapped onto a cartoon of a confined PC3 tumor cell–T cell dyad to emphasize compartment sidedness and communication mode. Tumor-associated responses include stress/metabolic and interferon-linked programs, antigen presentation, adhesion/ECM remodeling, and chemokine production, while T cell–associated responses include immediate-early activation genes, immune synapse components, MAPK feedback, and cytokine/chemokine secretion. Arrows highlight coordinated, bidirectional modules observed across dyads (e.g., a coupled “chemokine bloom”) and early activation dynamics captured within the defined 4.5 hour interaction window.

## References

1. Armingol, E., Baghdassarian, H. M. & Lewis, N. E. The diversification of methods for studying cell–cell interactions and communication. Nat. Rev. Genet. 25, 381–400 (2024).

2. Rood, J. E., Maartens, A., Hupalowska, A., Teichmann, S. A. & Regev, A. Impact of the Human Cell Atlas on medicine. Nat. Med. 28, 2486–2496 (2022).

3. Jerby-Arnon, L. & Regev, A. DIALOGUE maps multicellular programs in tissue from single-cell or spatial transcriptomics data. Nat. Biotechnol. 40, 1467–1477 (2022).

4. Jin, S. et al. Inference and analysis of cell-cell communication using CellChat. Nat. Commun. 12, 1088 (2021).

5. Larsson, L., Frisén, J. & Lundeberg, J. Spatially resolved transcriptomics adds a new dimension to genomics. Nat. Methods 18, 15–18 (2021).

6. Langerman, J. et al. Linking single-cell transcriptomes with secretion using SEC-seq. Nat. Protoc. 20, 2034–2055 (2025).

7. de Rutte, J. et al. Suspendable hydrogel Nanovials for massively parallel single-cell functional analysis and sorting. ACS Nano 16, 7242–7257 (2022).

8. Udani, S. et al. Associating growth factor secretions and transcriptomes of single cells in nanovials using SEC-seq. Nat. Nanotechnol. 19, 354–363 (2024).

9. Mao, Z. et al. Physical and in silico immunopeptidomic profiling of a cancer antigen prostatic acid phosphatase reveals targets enabling TCR isolation. Proc. Natl. Acad. Sci. 119, e2203410119 (2022).

10. Chen, X. et al. Overcoming T cell tolerance to tumor self-antigens through catch-bond engineering. Sci. Press (2025).

11. McGinnis, C. S., Murrow, L. M. & Gartner, Z. J. DoubletFinder: Doublet Detection in Single-Cell RNA Sequencing Data Using Artificial Nearest Neighbors. Cell Syst. 8, 329–337.e4 (2019).

12. Limoges, M.-A., Cloutier, M., Nandi, M., Ilangumaran, S. & Ramanathan, S. The GIMAP Family Proteins: An Incomplete Puzzle. Front. Immunol. Volume 12-2021, (2021).

13. Kobayashi, N., Takata, H., Yokota, S. & Takiguchi, M. Down-regulation of CXCR4 expression on human CD8+ T cells during peripheral differentiation. Eur. J. Immunol. 34, 3370–3378 (2004).

14. Arlt, A. & Schäfer, H. Role of the immediate early response 3 (IER3) gene in cellular stress response, inflammation and tumorigenesis. SFB 415 Specif. Pathophysiol. Signal Transduct. Pathw. 90, 545–552 (2011).

15. van Elsas, M. J. et al. Immunotherapy-activated T cells recruit and skew late-stage activated M1-like macrophages that are critical for therapeutic efficacy. Cancer Cell 42, 1032–1050.e10 (2024).

16. Cogswell, P. C., Mayo, M. W. & Baldwin, A. S., Jr. Involvement of Egr-1/RelA Synergy in Distinguishing T Cell Activation from Tumor Necrosis Factor-α–induced NF-κB1 Transcription. J. Exp. Med. 185, 491–498 (1997).

17. Wu, T., Womersley, H. J., Jiehao Ray, W., Scolnick, J. & Lih Feng, C. Time-resolved assessment of single-cell protein secretion by sequencing. Nat. Methods 20, 723–734 (2023).

18. Baatz, F. et al. Targeting BCL11B in CAR-engineered lymphoid progenitors drives NK-like cell development with prolonged anti-leukemic activity. Mol. Ther. 33, 1584–1607 (2025).

19. Forkel, H. et al. BCL11B depletion induces the development of highly cytotoxic innate T cells out of IL-15 stimulated peripheral blood αβ CD8+ T cells. OncoImmunology 11, 2148850 (2022).

20. Danaher, P. et al. Gene expression markers of Tumor Infiltrating Leukocytes. J. Immunother. Cancer 5, 18 (2017).

21. Binder, C. et al. CD2 Immunobiology. Front. Immunol. Volume 11-2020, (2020).

22. Kesarwani, P., Murali, A. K., Al-Khami, A. A. & Mehrotra, S. Redox Regulation of T-Cell Function: From Molecular Mechanisms to Significance in Human Health and Disease. Antioxid. Redox Signal. 18, 1497–1534 (2013).

23. Dunlock, V.-M. E. et al. Tetraspanin CD53 controls T cell immunity through regulation of CD45RO stability, mobility, and function. Cell Rep. 39, 111006 (2022).

24. Efremova, M., Vento-Tormo, M., Teichmann, S. A. & Vento-Tormo, R. CellPhoneDB: inferring cell–cell communication from combined expression of multi-subunit ligand–receptor complexes. Nat. Protoc. 15, 1484–1506 (2020).

25. Giladi, A. et al. Dissecting cellular crosstalk by sequencing physically interacting cells. Nat. Biotechnol. 38, 629–637 (2020).

26. Vonficht, D. et al. Ultra-high-scale cytometry-based cellular interaction mapping. Nat. Methods 22, 1887–1899 (2025).

27. Odagiu, L. et al. Early programming of CD8+ T cell response by the orphan nuclear receptor NR4A3. Proc. Natl. Acad. Sci. 117, 24392–24402 (2020).

28. Shao, H., Kono, D. H., Chen, L.-Y., Rubin, E. M. & Kaye, J. Induction of the Early Growth Response (Egr) Family of Transcription Factors during Thymic Selection. J. Exp. Med. 185, 731–744 (1997).

29. Elliot, T. A. E. et al. Antigen and checkpoint receptor engagement recalibrates T cell receptor signal strength. Immunity 54, 2481–2496.e6 (2021).

30. Chu, H. Y. et al. Dickkopf-1: A Promising Target for Cancer Immunotherapy. Front. Immunol. Volume 12-2021, (2021).

31. Yuan, C. R. & Lee, T. K. W. TM4SF1 – A new immune target for treatment of hepatocellular carcinoma: Editorial on “Targeting TM4SF1 promotes tumor senescence enhancing CD8+ T cell cytotoxic function in hepatocellular carcinoma”. CMH 31, 646–649 (2025).

32. Cheng, R. Y.-H. et al. SEC-seq: association of molecular signatures with antibody secretion in thousands of single human plasma cells. Nat. Commun. 14, (2023).

33. Koo, D. et al. Defining T cell receptor repertoires using nanovial-based binding and functional screening. Proc. Natl. Acad. Sci. 121, (2024).

34. Koo, D. et al. Optimizing cell therapy by sorting cells with high extracellular vesicle secretion. Nat. Commun. 15, 4870 (2024).

35. Dura, B. et al. Profiling lymphocyte interactions at the single-cell level by microfluidic cell pairing. Nat. Commun. 6, 5940 (2015).

36. Shaik, F. A. et al. Pairing cells of different sizes in a microfluidic device for immunological synapse monitoring. Lab. Chip 22, 908–920 (2022).

37. Frimat, J.-P. et al. A microfluidic array with cellular valving for single cell co-culture. Lab. Chip 11, 231–237 (2011).

38. Şen, M., Ino, K., Ramón-Azcón, J., Shiku, H. & Matsue, T. Cell pairing using a dielectrophoresis-based device with interdigitated array electrodes. Lab. Chip 13, 3650–3652 (2013).

39. Mao, T. et al. Single-Target Pairing System (StarPair) for Large-Scale Interrogation of Cell–Cell Interactions. Adv. Sci. n/a, e13951 (2025).

40. Challa, D. et al. Function-first discovery of high affinity monoclonal antibodies using Nanovial-based plasma B cell screening. bioRxiv 2024.08.15.608174 (2024) doi:10.1101/2024.08.15.608174.

41. Stoeckius, M. et al. Simultaneous epitope and transcriptome measurement in single cells. Nat. Methods 14, 865–868 (2017).

41. Chromatin accessibility profiling methods. Nat. Rev. Methods Primer 1, 11 (2021).

43. Bartosovic, M., Kabbe, M. & Castelo-Branco, G. Single-cell CUT&Tag profiles histone modifications and transcription factors in complex tissues. Nat. Biotechnol. 39, 825–835 (2021).

